# Distances and charges along the Orai1 nexus-TM3 interface control STIM1-binding and pore opening

**DOI:** 10.1101/2025.08.08.669298

**Authors:** Julia Söllner, Magdalena Prantl, Hadil Najjar, Veronika Aichner, Ana Marija Suta, Yuliia Nazarenko, Selina Haranth, Maximilian Fröhlich, Diana Thallinger, Tamara Radiskovic, Christopher Mairhofer, William Nzegge, Sarah Weiß, Heinrich Krobath, Mario Waser, Isabella Derler

**Affiliations:** Institute of Biophysics, JKU Life Science Center, Johannes Kepler University Linz, Gruberstraße 40, 4020 Linz, Austria; Institute of Theoretical Physics, Johannes Kepler University Linz, Altenbergerstraße 69, 4040 Linz, Austria; Institute of Organic Chemistry, Johannes Kepler University Linz, Altenbergerstraße 69, 4040 Linz, Austria

## Abstract

Calcium (Ca^2+^) influx through the Ca^2+^ release-activated Ca^2+^ (CRAC) channel is triggered by binding of the Ca^2+^ sensor Stromal Interaction Molecule 1 (STIM1) to the pore-forming Orai1 complex, primarily to its cytosolic C-termini. These C-termini connect to the transmembrane domain (TM) 4 via the flexible nexus region, proposed to transmit the activation signal from the STIM1-binding site to the central pore via concentrically arranged TM domains. However, the conformational dynamics of the nexus-TM3 interface required for channel gating remain elusive. Here, we investigate its role using unnatural amino acid (UAA)-based photo- and chemical crosslinking at individual positions within the nexus-TM3 interface combined with conventional site-directed mutagenesis. We report that a widening of the nexus-TM3 interface is essential for STIM1-mediated pore opening, while hydrophobicity and contact distances in the upper nexus-TM3 interface fine-tune signal propagation to the pore. These findings underscore the relevance of the nexus-TM3 dynamics for proper Orai1 function.

## Introduction

Calcium ions (Ca²⁺) are vital second messengers in numerous signaling pathways that regulate key cellular processes such as gene expression, muscle contraction, immune response, and cell proliferation ^1–3^. One of the most prominent Ca²⁺ entry routes is the Ca²⁺ release-activated Ca²⁺ (CRAC) channel, which plays a crucial role especially in immune cells. It consists of the pore-forming protein Orai1 in the plasma membrane (PM) and the Ca²⁺ sensor Stromal Interaction Molecule 1 (STIM1) in the membrane of the endoplasmic reticulum (ER), the main Ca^2+^ store of a cell ^4–7^. Upon ER Ca²⁺ store-depletion, STIM1 undergoes a conformational rearrangement, oligomerizes, and translocates to ER–PM junctions where it binds to and activates Orai1, thereby triggering Ca²⁺ influx ^8–10^. Several mutations in either STIM1 or Orai1 can lead to loss- or gain-of-function (L-/GoF) of the CRAC channel complex. This can cause diseases such as severe immunodeficiencies, Stormorken-like syndrome or tubular aggregate myopathy, highlighting the physiological relevance and the need for a deeper understanding of the molecular mechanisms of the tightly controlled CRAC channel activation ^11–13^.

In this work, we focus on the human Orai1 (hOrai1) isoform denoted as Orai1. It forms a unique hexameric complex, whereof each monomer comprises four transmembrane (TM) helices, two extracellular and one intracellular loop, and a cytosolic N- and C-terminus. The six TM1 helices form the highly Ca²⁺ selective pore, which is enclosed by a ring of the six TM2-TM3 helices and TM4 in the periphery ^14–17^. Pore opening is initiated by STIM1-binding to its main coupling site, the C-terminus of Orai1, triggering a series of conformational changes across the entire channel, resulting in pore opening ^12, 18–22^. The Orai1 C-terminus is connected via the so-called nexus region to TM4, which has been reported to be important for pore opening ^23^. The flexible nexus region consists of two segments – the lower flexible nexus (K265, H264, S263) and the upper hydrophobic nexus (V262, L261) (Fig. 1a). It is highly conserved in *Drosophila melanogaster* Orai (dOrai) and a structural comparison between the closed and open dOrai crystal structures suggests that the kinked, antiparallel C-termini in the resting conformation unlatch and straighten upon channel activation (Fig. 1a) ^14, 16^. While such conformational changes appear less pronounced in the corresponding cryo-EM structures (PDB: 7KR5)^15^, cysteine cross-linking studies on the Orai1 C-terminus again indicate its conformational change upon STIM1-coupling ^24^. A recent study by Zhou et al.^23^ proposes that STIM1-binding provokes a bending of the lower flexible nexus, which propagates to the upper nexus-TM3 interface to induce pore opening. We recently published that non-pore-lining TM interfaces, which are connected to the nexus region at the cytosolic side, undergo STIM1-induced widening crucial for pore opening^25^. However, the exact conformational changes and the role of the nexus-TM3 interface in the physiologically relevant STIM1-mediated pore opening process still remain unclear.

**Figure 1:**
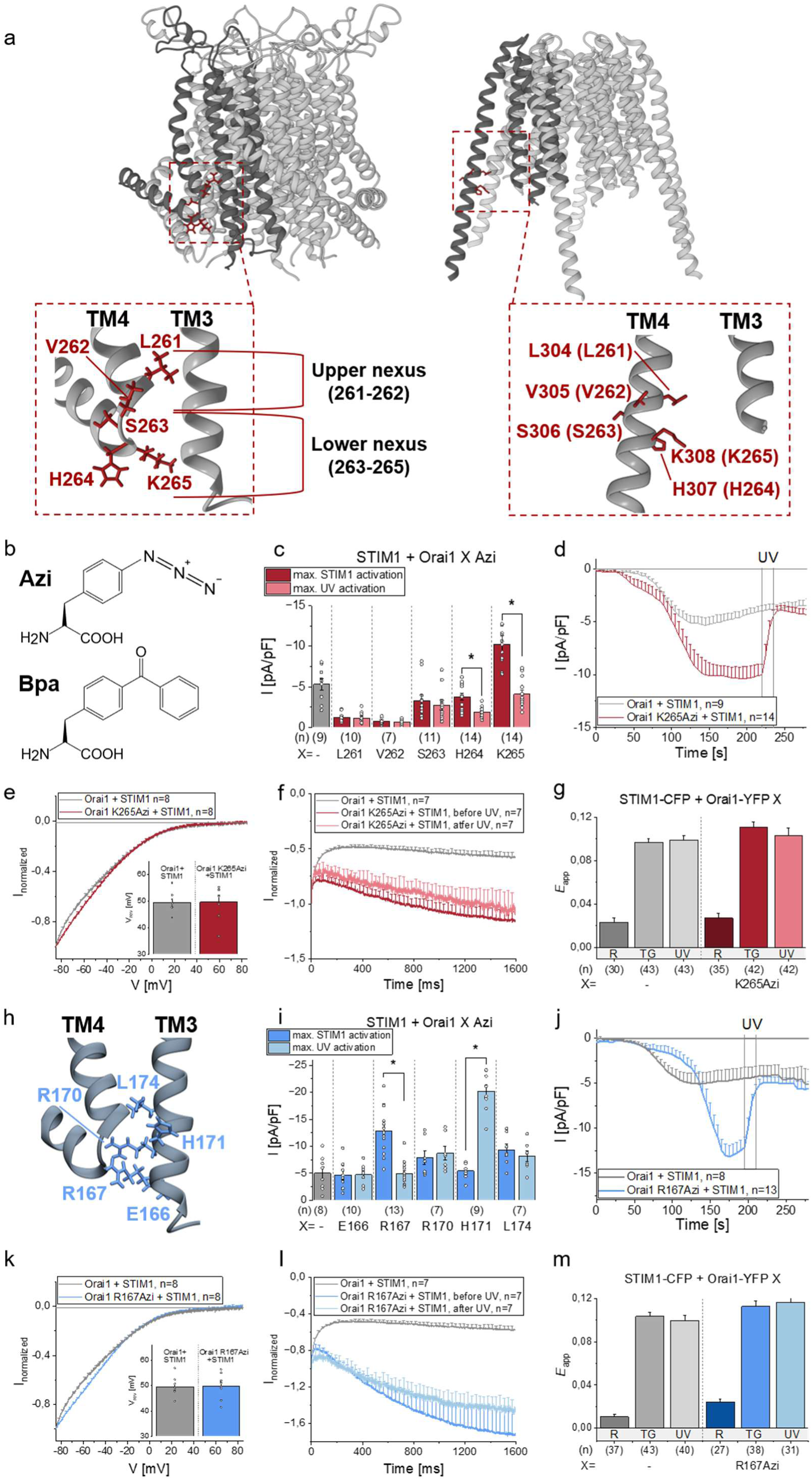
Photocrosslinking UAAs at sites in the nexus and opposite in TM3 allow upon STIM1-mediated activation UV-induced current reduction. **a)** Scheme showing closed hOrai1 (left; PDB: 4HKR) and open dOrai (right; PDB: 6BBF) structures with insets depicting the upper and lower nexus region (LVSHK; 261-265 in hOrai1/ 304-307 in dOrai) with tested residues highlighted in red. Numbers indicate positions in hOrai1 or dOrai with those of hOrai1 in brackets. b) Chemical structures of Azi and Bpa. c) i) Bar diagrams showing maximal current densities after STIM1-activation and 15s UV-illumination of c) Orai1 nexus-mutants and i) Orai1 TM3-mutants containing Azi. Statistical significance before and after UV-illumination are indicated by asterisk * (p< 0.05). d) j) Time course of current densities after whole cell break-in of Orai1 WT and d) Orai1 K265Azi and j) Orai1 R167Azi co-expressed with STIM1. UV-light is applied for 15s (t= 220s (d)/195s (j)), after maximum STIM1-mediated currents have been reached. e) k) Normalized I/V relationships of Orai1 WT and e) Orai1 K265Azi and k) Orai1 R167Azi in the presence of STIM1. Inlet represents the reversal potential (Vrev) of STIM1-mediated currents. f) l) Time course showing normalized currents of Orai1 WT compared to f) Orai1 K265Azi and l) Orai1 R167Azi before and after 15s UV-illumination in the presence of STIM1. Currents were obtained from the application of a voltage step of -110 mV. g) m) Bar diagrams exhibiting FRET efficiency (Eapp) detecting the interaction of STIM1-CFP with Orai1-YFP WT compared to g) Orai1-YFP K265Azi and m) Orai1-YFP R167Azi at resting conditions (R), after store-depletion with 1 µM Thapsigargin (TG) and after 60s UV-illumination (UV). h) Schematic representation of the nexus-TM3 interface highlighting tested residues in TM3. Data represent mean values ± SEM of indicated number (n) of experiments. Single values are indicated as grey circles. Detailed statistic values are shown in Supplementary Table 1.

To uncover the molecular mechanism and essential conformational rearrangements within the Orai1 channel complex, we successfully demonstrated that the emerging genetic code expansion (GCE) technology is a powerful tool ^25, 26^. Thereby, an orthogonal aminoacyl-tRNA synthetase/tRNA pair is introduced into the host organism, enabling the incorporation of a certain unnatural amino acid (UAA) at a site-specifically introduced amber stop codon (TAG) during the native ribosomal translation ^27, 28^.

In this study, we used this technique to introduce UAAs with photo- and chemical crosslinking capabilities into the Orai1 nexus-TM3 interface to gain, combined with functional studies, a deeper and more precise understanding of conformational dynamics within this region during channel opening. We seek to uncover another missing piece of the puzzle in understanding the molecular mechanisms by which the STIM1-activation signal is transmitted from the peripheral C-terminus, through the nexus, to the entire channel complex and the central pore of Orai1.

## Results

### Photocrosslinking UAAs at sites in the nexus and opposite in TM3 allow upon STIM1-mediated activation UV-induced current reduction

To shed light on the exact conformational dynamics within the nexus and its interactions with TM3 critical for STIM1-mediated activation, we incorporated photocrosslinking UAAs, specifically p-azido-L-phenylalanine (Azi) and p-benzoyl-L-phenylalanine (Bpa) (Fig. 1b), both capable of forming a covalent bond with any residue within 3-4 Å upon UV-light exposure^29, 30^, within this region.

An initial mutational screen with photocrosslinking UAAs, Azi or Bpa, through the nexus region (^261^LVSHK^265^) revealed one outstanding site, K265. Insertion of Azi at this position (Orai1 K265Azi) leads to significantly increased Orai1 wild-type (WT) currents upon STIM1 co-expression before exposure to UV-light, which were significantly reduced upon UV-illumination to current levels under WT conditions (Fig. 1c,d). In a negative control experiment, the Orai1 C-terminal mutation L273D, that impairs STIM1-binding and STIM1-mediated activation^31^, also blocked STIM1-induced activation of Orai1 K265Azi L273D (Supp. Fig. 1a). The current/voltage (I/V) relationship of STIM1 + Orai1 K265Azi currents showed a comparable robust inward rectification and a reversal potential of approximately +50 mV like STIM1-mediated Orai1 WT currents (Fig. 1e). Fast Ca^2+^ dependent inactivation (FCDI) was reduced compared to that of STIM1-mediated Orai1 WT currents and was followed by a reactivation phase (Fig. 1f), which might be related to the strongly increased store-operated currents. To investigate whether the elevated store-operated currents of Orai1 K265Azi or the reduction in current size after UV-light exposure were due to changes in interactions with STIM1, fluorescence resonance energy transfer (FRET) of Orai1-YFP K265Azi with STIM1-CFP compared to WT conditions was measured before and after store-depletion via Thapsigargin (TG) and following UV-exposure. No significant changes in the FRET efficiency were detected using these different conditions (Fig. 1g, Supp. Fig. 1b), which indicates comparable STIM1-binding to Orai1 WT and Orai1 K265Azi.

In contrast to Orai1 K265Azi, Azi substitutions of other residues along the nexus led to smaller STIM1-mediated currents compared to Orai1 WT. Orai1 H264Azi showed a significant UV-mediated current reduction following maximal STIM1-induced activation and for Orai1 S263Azi, a similar tendency was observed (Fig. 1c). Among the corresponding Bpa-nexus mutants, also Orai1 K265Bpa tended to exhibit enhanced currents compared to Orai1 WT, which reduced slightly after UV-exposure, whereas other Bpa-nexus mutants showed only weak STIM1-mediated activation, which did not alter after UV-illumination (Supp. Fig. 1c).

As several side chains in TM3 point directly towards the nexus region (Fig. 1h), we performed a second screen by incorporating Azi/ Bpa at positions in TM3 which are located opposite to the nexus. Here, Orai1 R167Azi showed significantly enhanced STIM1-mediated currents with UV-induced current reduction to WT-like levels (Fig. 1i,j). The I/V relationship was like WT (Fig. 1k), whereas FCDI occurring within 50ms, was less pronounced compared to Orai1 WT and was followed by a reactivation phase (Fig. 1l). The binding of Orai1-YFP R167Azi and STIM1-CFP appears unaltered as indicated by FRET experiments (Fig. 1m, Supp. Fig. 1b). Upon Bpa incorporation, a similar, but not significant trend was detected for Orai1 R167Bpa (Supp. Fig. 1d).

Towards the upper nexus, the incorporation of Azi but not Bpa at Orai1 H171 resulted in WT-like currents, that were drastically increased upon UV-illumination (Fig. 1i, Supp. Fig. 1d). A similar UV-induced current enhancement yet upon introducing Bpa rather than Azi was observed for Orai1 L174Bpa, as published in our previous study^26^ (Fig. 1i, Supp. Fig. 1d). Other corresponding Azi- or Bpa-mutants along the upper nexus exhibited STIM1-mediated activation similar to Orai1 WT and were unaffected by UV-light exposure.

Our UAA-containing mutants showed neither constitutive activity nor UV-induced current alterations in the absence of STIM1 as shown for all Azi-mutants and selected Bpa-mutants (Supp. Fig. 1e,f). Only Orai1 L174Bpa exhibits UV-induced activation already in the absence of STIM1 (Fig. 1f), as already discovered in our previous study ^26^.

Summarizing, UV-exposure upon UAA-incorporation in and opposite to the lower nexus region (K265, R167) results in strong STIM1-mediated currents and UV-induced current reduction. In contrast, UV-illumination of UAAs in TM3 facing the upper nexus region (H171, L174) leads to robust STIM1-mediated current increases.

### Aromatic substitutions at K265 increase STIM1-mediated currents of Orai1

Since for both positions, K265 and R167 (Fig. 2a), in particular the incorporation of the bulky Azi, resulted in significantly enhanced STIM1-mediated currents, we hypothesized that a widening of the nexus-TM3 interface due to the increased size of the inserted UAA, could be responsible for the enhanced current size. We further speculate that the UV-mediated current reduction is potentially associated with a nexus-TM3 distance dependence. To probe for a relation between distance and current size, we mutated K265 and R167 to bulky aromatic as well as small amino acids for negative control (K265A/F/Y/W; R167A/F/Y/W). Indeed, electrophysiological experiments revealed significantly enhanced STIM1-mediated currents for Orai1 K265F/Y compared to Orai1 WT but not for Orai1 K265A (Fig. 2b,c). Despite the greatly increased currents, the I/V-relationship of STIM1-mediated currents was similar to Orai1 WT, as exemplarily shown for Orai1 K265Y (Fig. 2d). Orai1 K265W, similar to Orai1 K265Bpa, showed a tendency for, but not significantly increased currents compared to WT conditions (Fig. 2c), probably due to steric interference of the bulky tryptophan or the formation of inhibitory hydrophobic interactions.

**Figure 2:**
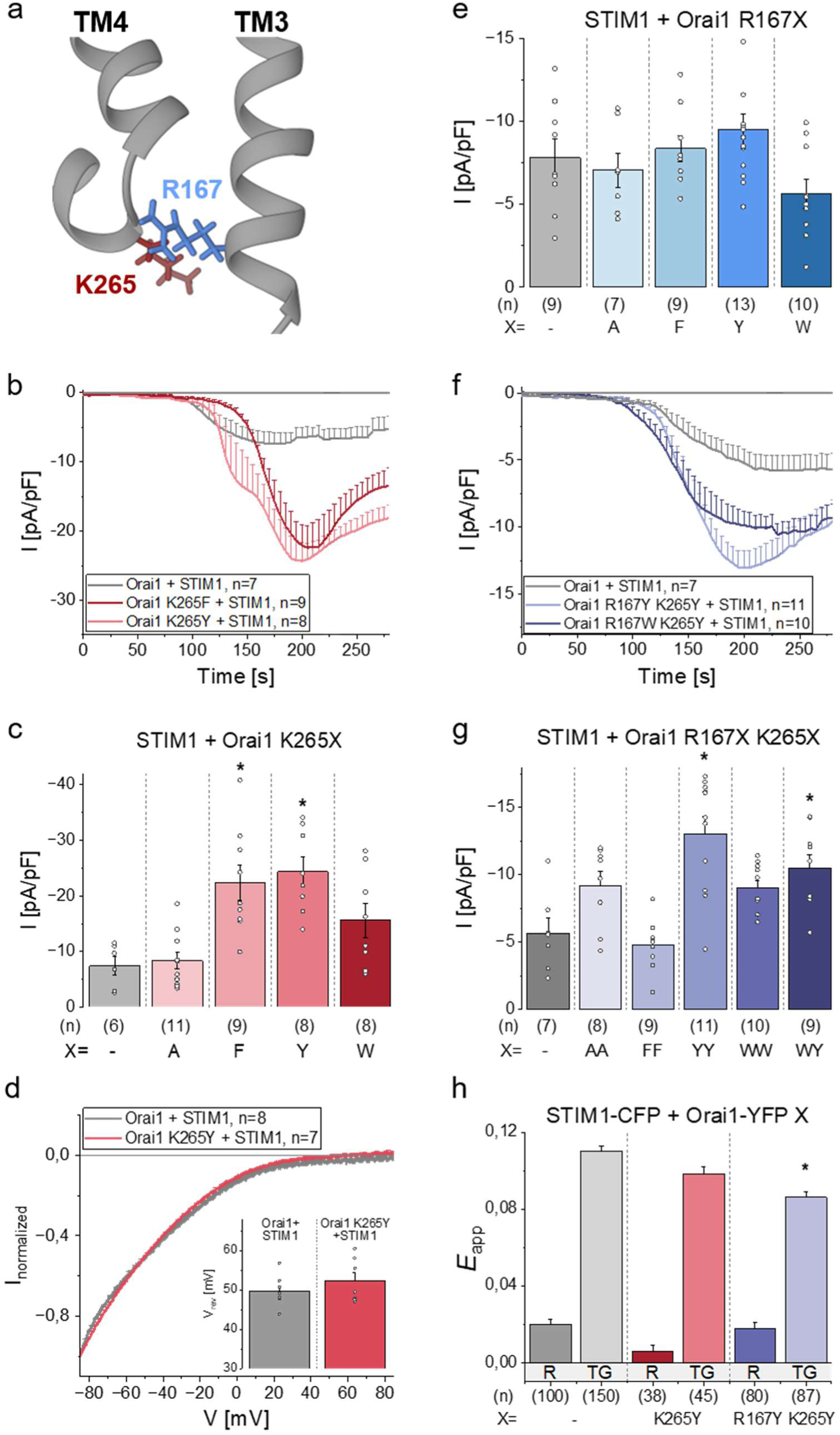
Aromatic substitutions at K265 increase STIM1-mediated currents of Orai1. **a)** Schematic representation of key residues K265 and R167 in the nexus-TM3 interface of Orai1. **b) f)** Time course of current densities after whole cell break-in of Orai1 WT and b) Orai1 K265F/Y and f) Orai1 R167Y K265Y and Orai1 R167W K265Y in the presence of STIM1. **c) e) g)** Bar diagrams showing maximal current densities after STIM1-activation of Orai1 WT and c) Orai1 K265A/F/Y/W, e) Orai1 R167A/F/Y/W and g) Orai1 R167A K265A, Orai1 R167F K265F, Orai1 R167W K265W and Orai1 R167W K265Y. Statistical significance of Orai1 mutants compared to Orai1 WT are indicated by asterisk **\*** (p< 0.05). **d)** Normalized I/V relationships of Orai1 WT and Orai1 K265Y in the presence of STIM1. Inlet represents the reversal potential (Vrev) of STIM1-activated Orai1 currents. **h)** Bar diagrams exhibiting the FRET efficiency (Eapp) detecting the interaction of STIM1-CFP with Orai1-YFP WT compared to Orai1-YFP K265Y and Orai1-YFP R167Y K265Y at resting conditions (R) and after store-depletion with 1 µM Thapsigargin (TG). Statistical significance of Orai1 mutants compared to Orai1 WT at similar conditions are indicated as asterisk **\*** (p< 0.05). Single values are indicated as grey circles. Data represent mean values ± SEM of indicated number (n) of experiments. Detailed statistic values are shown in Supplementary Table 1.

In contrast, maximum WT-like STIM1-mediated currents were observed for Orai1 R167 mutants, regardless of the size of the incorporated amino acid (Fig. 2e). Next, we introduced double point mutations (R167A/F/Y/W K265A/F/Y/W) to test whether the current could be enhanced. Only for the mutants, Orai1 R167Y K265Y and Orai1 R167W K265Y, STIM1-mediated currents were significantly increased compared to WT, albeit not reaching the level of the Orai1 K265Y single mutant (Fig. 2f,g). The FRET signal was significantly lower for Orai1 R167Y K265Y, but not for Orai1 K265Y compared to Orai1 WT, indicating altered STIM1-binding for the double-Y mutant (Fig. 2h). Interestingly, the current size of Orai1 R167F K265F was markedly lower than that of the single mutant Orai1 K265F (Fig. 2c,g), possibly due to local formation of inhibitory hydrophobic interactions, which might reflect in reduced backbone flexibility and the suppression of the propagation of the opening signal.

In summary, increasing the amino acid side chain size at position K265 but not at position R167, both situated in the nexus-TM3 interface, increases STIM1-mediated currents. Some double aromatic substitutions at these positions can enhance currents but also impair STIM1-coupling compared to Orai1 WT.

### Alanine substitutions in TM3 opposite the UAA in the nexus leave UV-mediated current decrease unaffected, but, in some cases, decrease STIM1-induced maximal activation

Since bulky aromatic side chains along the nexus trigger significant enhancements of STIM1-mediated currents, we hypothesized that the observed decrease in STIM1-induced currents after UV-light application could be due to a decrease in the distance of the nexus region caused by photocrosslinking. We therefore tested whether a reduction in side-chain sizes of residues at critical positions in TM3 opposite to a UAA in the nexus region could counteract UV-induced current reduction. Hence, we combined Orai1 K265Azi with mutations in TM3 (E166, R167, R170) to alanine (Fig. 3a). However, for all double or triple mutants, UV-illumination still resulted in current decrease after maximal STIM1-induced activation before UV-light exposure (Fig. 3b, Supp. Fig. 2a). Interestingly, the combination of K265Azi with E166A or in some cases R170A revealed approximately 50% lower STIM1-mediated maximal currents compared to Orai1 K265Azi alone. Notably, double or triple mutants showing similar maximal currents like Orai1 K265Azi exhibited UV-induced current reduction to levels 50% reduced compared to WT conditions. In contrast, mutants exhibiting reduced maximal STIM1-mediated currents compared to Orai1 K265Azi, revealed UV-mediated current decrease even 75% reduced compared to those under WT conditions. Overall, this might be related to the reduced side chain volume. FRET levels were also significantly reduced for Orai1-YFP K265Azi E166A and Orai1-YFP K265Azi E166A R167A/R170A compared to Orai1 K265Azi and Orai1 WT, all co-expressed with STIM1-CFP (Fig. 3c). Hence, STIM1-binding to these double and triple mutants might be affected, potentially acting as another cause for decreased current levels. Thus, reducing amino acid side chain size of oppositely located residues does not eliminate UV-induced de-activation, but in some cases can lead to overall lower current density levels than for STIM1-activated Orai1.

**Figure 3:**
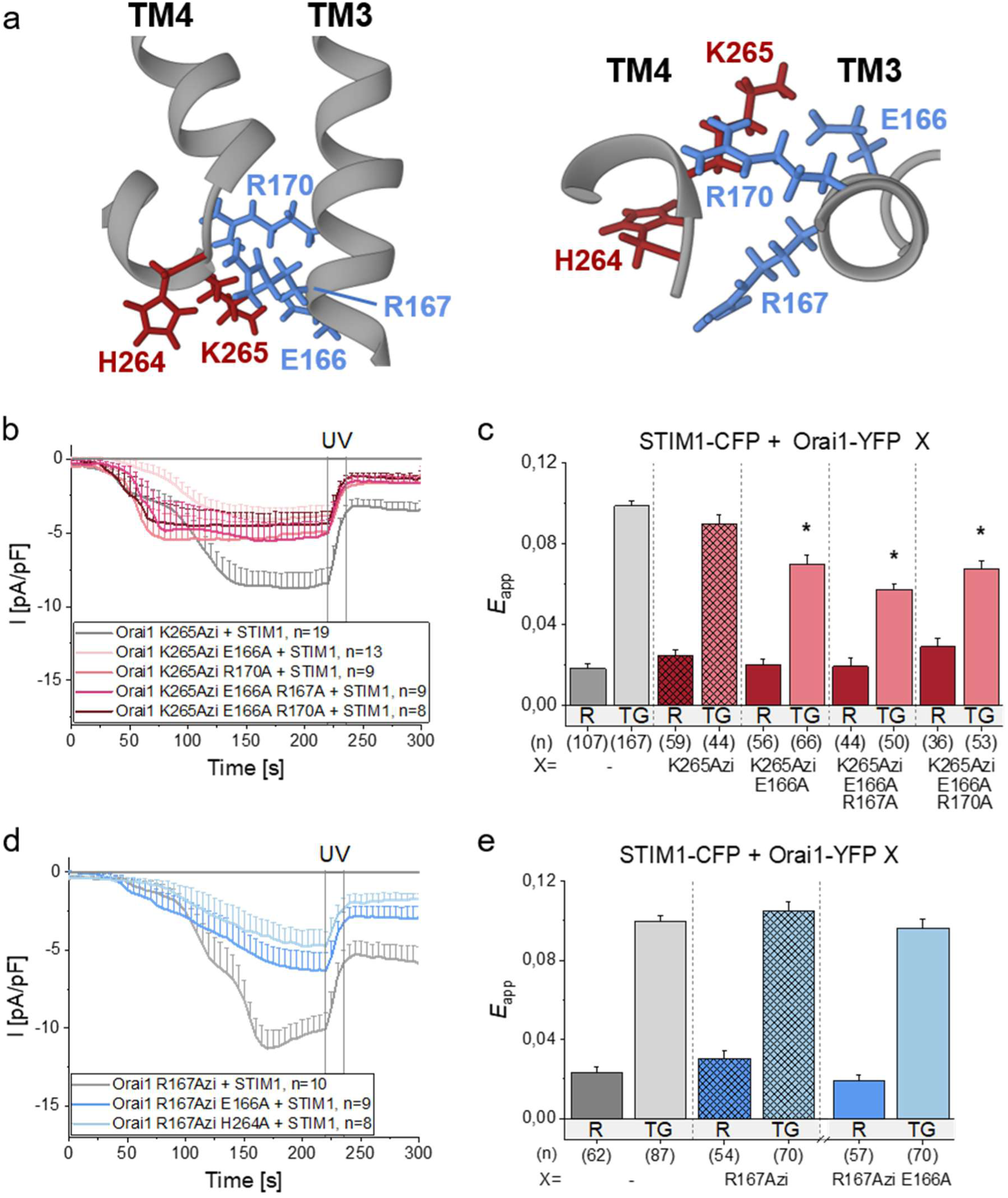
Alanine substitutions in TM3 opposite the UAA in the nexus leave UV-mediated current decrease unaffected, but, in some cases, decrease STIM1-induced maximal activation. **a)** Schematic representation of the side view (left) and top view (right) of the Orai1 nexus-TM3 interface highlighting key residues. **b) d)** Time course of current densities after whole cell break-in of b) Orai1 K265Azi and Orai1 K265Azi E166A/R170A/E166A R167A/E166A R170A and d) Orai1 R167Azi and Orai1 R167Azi E166A/H264A in the presence of STIM1. UV-light is applied for 15s at t= 220s, after maximum STIM1-mediated currents were reached. **c) e)** Bar diagrams showing FRET efficiency (Eapp) detecting the binding interaction of STIM1-CFP with Orai1-YFP WT and c) Orai1-YFP K265Azi/ K265Azi E166A/E166A R167A/E166A R170A and e) Orai1-YFP R167Azi/R167Azi E166A at resting conditions (R) and after store-depletion with 1 µM Thapsigargin (TG). Statistical significance of Orai1 mutants compared to Orai1 WT at similar conditions are indicated by asterisk **\*** (p< 0.05). Data represent mean values ± SEM of indicated number (n) of experiments. Detailed statistic values are shown in Supplementary Table 1.

A similar approach was performed for Orai1 R167Azi, in which different residues close to or opposite R167 (E166, R170, H264, K265) were mutated again to an alanine (Fig. 3a). Analogous to mutant Orai1 K265Azi, all tested Orai1 R167Azi-alanine-mutants showed STIM1-mediated activation, which reduced after UV-exposure (Supp. Fig. 2b). Interestingly, the combination of R167Azi with E166A or with H264A resulted in lower STIM1-mediated currents followed by further UV-induced current reductions below levels reached under WT conditions (Fig. 3d). While FRET levels for the double mutant Orai1 R167Azi E166A were similar to R167Azi under resting conditions as well as after store-depletion (Fig. 3e). This highlights again a potential role of side chain size at the nexus-TM3 interface in STIM1-induced Orai1 activation.

In conclusion, decreasing side chain size in the surroundings of Orai1 K265Azi or Orai1 R167Azi is not sufficient to eliminate the UV-induced reduction of STIM1-mediated currents. Nevertheless, in combination with E166A and R170A or H264A, respectively, both K265Azi and R167Azi showed strongly decreased STIM1-mediated current levels, indicating an important function of their local interaction partners in the nexus-TM3 interface in pore opening and/or signal transduction.

### Combination of K265Azi/Bpa or R167Azi/Bpa with mutations triggering constitutive activity eliminates UV-induced current decrease

Next, we investigated the effect of combining K265Azi or R167Azi with mutations within Orai1 which alone led to constitutive Ca^2+^ influx also under store-replete conditions. Here, we focused on several known GoF-mutations located across all four TM domains including V102A in TM1, H134A in TM2, V181K in TM3 and V181W A254W in TM3-TM4 (Fig. 4a). All of them lead to constitutively active Orai1 currents with almost CRAC channel-like I/V relationship and reversal potential^32, 33^ (Fröhlich et al., unpublished), except V102A resulting in non-selective constitutive currents ^34^. While V102A disrupts the hydrophobic gate in the pore^35^, H134A destabilizes the closed state by abolishing the function of the histidine as a steric brake ^36^. For V181K^37^ and V181W A254W (Fröhlich et al., unpublished), we reported that they affect the non-pore-lining TM interfaces, involving their widening, which in turn contributes to constitutive pore opening. Hence, it is known that V181K and V181W A254W affect the TM3-TM4 interface, while it remains unclear whether V102A and H134A also impact the peripheral channel segments. Therefore, we speculated that photocrosslinking of K265Azi impacts Orai1 V102A and Orai1 H134A differently than Orai1 V181K and Orai1 V181W A254W. The combination of the different GoF-mutations with K265Azi led in all cases to constitutive currents in the absence of STIM1 co-expression (Fig. 4b,c) with I/V relationships and reversal potentials comparable to previous reports, as exemplarily shown for Orai1 K265Azi V102A/H134A/V181K (Supp. Fig. 3a) ^32, 37, 38^. However, these mutation combinations eliminated the UV-induced current decrease (Fig. 4b,c). This indicates that the GoF-mutations trigger different conformational changes in the nexus-TM3 interface than STIM1, which hinder functional effects of UV-illumination. In the presence of STIM1, all double mutants exhibited constitutively active currents with I/V-relationships reaching a reversal potential of around +50 mV for Orai1 K265Azi V102A/H134A/V181K (Supp. Fig. 3b), consistent with previous studies ^32, 37, 38^. Interestingly, in the presence of STIM1, UV-induced photocrosslinking had a slight, but no significant effect on the tested Orai1 GoF-K265Azi mutants (Fig. 4d,e). STIM1-binding was not negatively affected as exemplarily shown for STIM1-CFP and Orai1 K265Azi V181K-YFP (Supp. Fig. 3c). Also, for Orai1 R167Azi V181K only a weak UV-induced current reduction was observed in the absence as well as in the presence of STIM1 (Fig. 4f).

**Figure 4:**
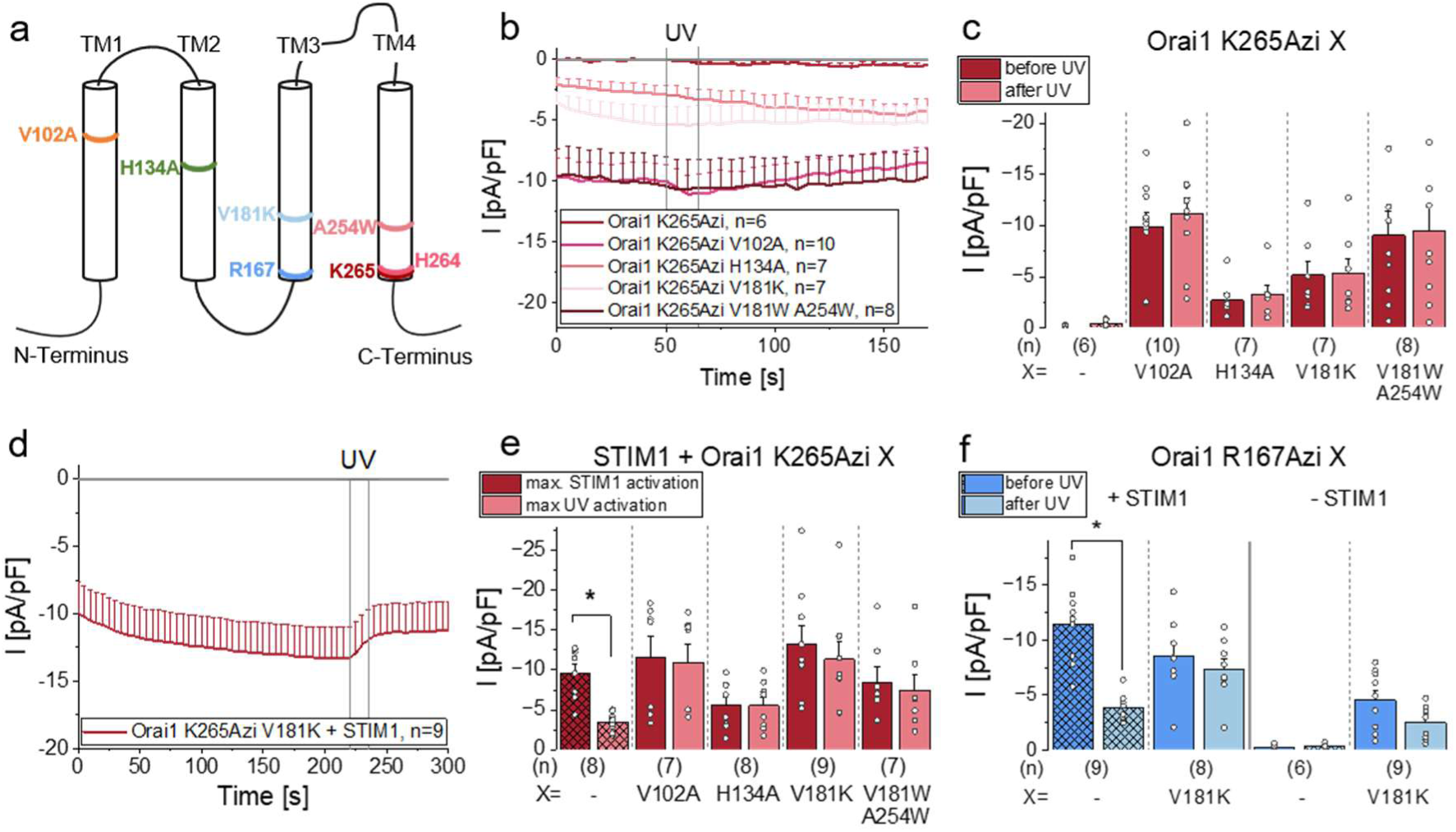
Combination of K265Azi/Bpa or R167Azi/Bpa with mutations triggering constitutive activity eliminates UV-induced current decrease. a) Schematic representation of the Orai1 subunit highlighting tested GoF-mutants in TM1-TM4. b) **d)** Time course of current densities after whole cell break-in of b) Orai1 K265Azi and Orai1 K265Azi V102A/H134A/V181K/V181W A254W in the absence of STIM1 and of d) Orai1 K265Azi V181K in the presence of STIM1. UV-light is applied for 15s at t= 50s (b)/ 220s (d). c) **e)** Bar diagrams exhibiting maximal current densities before and after 15s UV-illumination of Orai1 K265Azi and Orai1 K265Azi V102A/H134A/V181K/V181W A254W in the c) absence of STIM1 and e) presence of STIM1. Statistical significance before and after UV-illumination are indicated by asterisk **\*** (p< 0.05). **f)** Bar diagram showing maximal current densities after STIM1-activation and after 15s UV-illumination of Orai1 R167Azi and Orai1 R167Azi V181K in the presence (left) and absence (right) of STIM1. Statistical significance before and after UV-illumination are indicated by asterisk **\*** (p< 0.05). Single values are indicated as grey circles. Data represent mean values ± SEM of indicated number (n) of experiments. Detailed statistic values are shown in Supplementary Table 1.

Additionally tested Bpa-mutants (Orai1 K265Bpa V181K, Orai1 R167Bpa V181K) again exhibited no significant UV-mediated current decrease in the absence or presence of co-expressed STIM1 (Supp. Fig. 3d).

Furthermore, we investigated the nexus mutant Orai1 H264Azi in combination with GoF-mutants H134A and V181K. For both GoF-H264Azi mutants, constitutive activity but no significant effect of UV-light illumination on current size was observed, both in the absence and presence of STIM1 (Supp. Fig. 3e).

These results indicate that UV-induced photocrosslinking of K265Azi, H264Azi or R167Azi in the presence of the tested GoF-mutations is unable to induce conformational changes along the nexus-TM3 interface in the absence of STIM1. We hypothesize that, due to a structural change in this peripheral channel segment, such as putative widening of the non-pore-lining TM interface in the GoF-mutants, Azi at K265 and R167 can hardly reach its crosslinking partner. STIM1 seems to induce some structural changes at the nexus-TM3 interface which allow at least for the TM3-TM4 GoF-mutants slight, but not significant UV-induced functional changes due to potential structural rearrangements in this peripheral channel segment.

Taken together, UV-induced current decrease of Orai1 K265Azi, Orai1 H264Azi and Orai1 R167Azi is lost in combination with all tested GoF-mutations even in the presence of STIM1.

### Restraining the nexus-TM3 interface via photo- and chemical crosslinking impairs STIM1-mediated maximal activation

As we assume a widening along the nexus-TM3 interface during STIM1-binding and channel opening, we wondered whether exposure to UV-light prior to STIM1-binding might interfere with opening-relevant conformational changes to occur and thereby lower maximal current size. To this end, we compared the effects of two different experimental activation protocols on critical Orai1 UAA-mutants in the patch-clamp experiment: i) the initial application of UV-light immediately after entering the whole-cell configuration, hence, before STIM1 binds to the Orai1 mutant, followed by passive store-depletion-induced STIM1-mediated activation and ii) allowing for STIM1-binding and maximal STIM1-mediated currents to be reached and subsequent illumination of UV-light for photocrosslinking, as in previously shown measurements. Indeed, UV-exposure before STIM1-binding nearly abolished STIM1-mediated currents for Orai1 K265Azi. Similar results were detected for some K265Azi-double and triple mutants, as exemplarily shown for Orai1 K265Azi E166A and Orai1 K265Azi E166A R167A (Fig. 5a,b). Hence, the UV-induced photocrosslink of K265Azi hinders STIM1-mediated activation of the channel, probably by preventing conformational rearrangements of the nexus-TM3 interface, that might be necessary for proper pore opening. STIM1-binding itself is not impaired by applying UV-light prior to STIM1-binding, as FRET measurements showed similar results to Orai1 WT for both experimental conditions (Supp. Fig. 4a).

**Figure 5:**
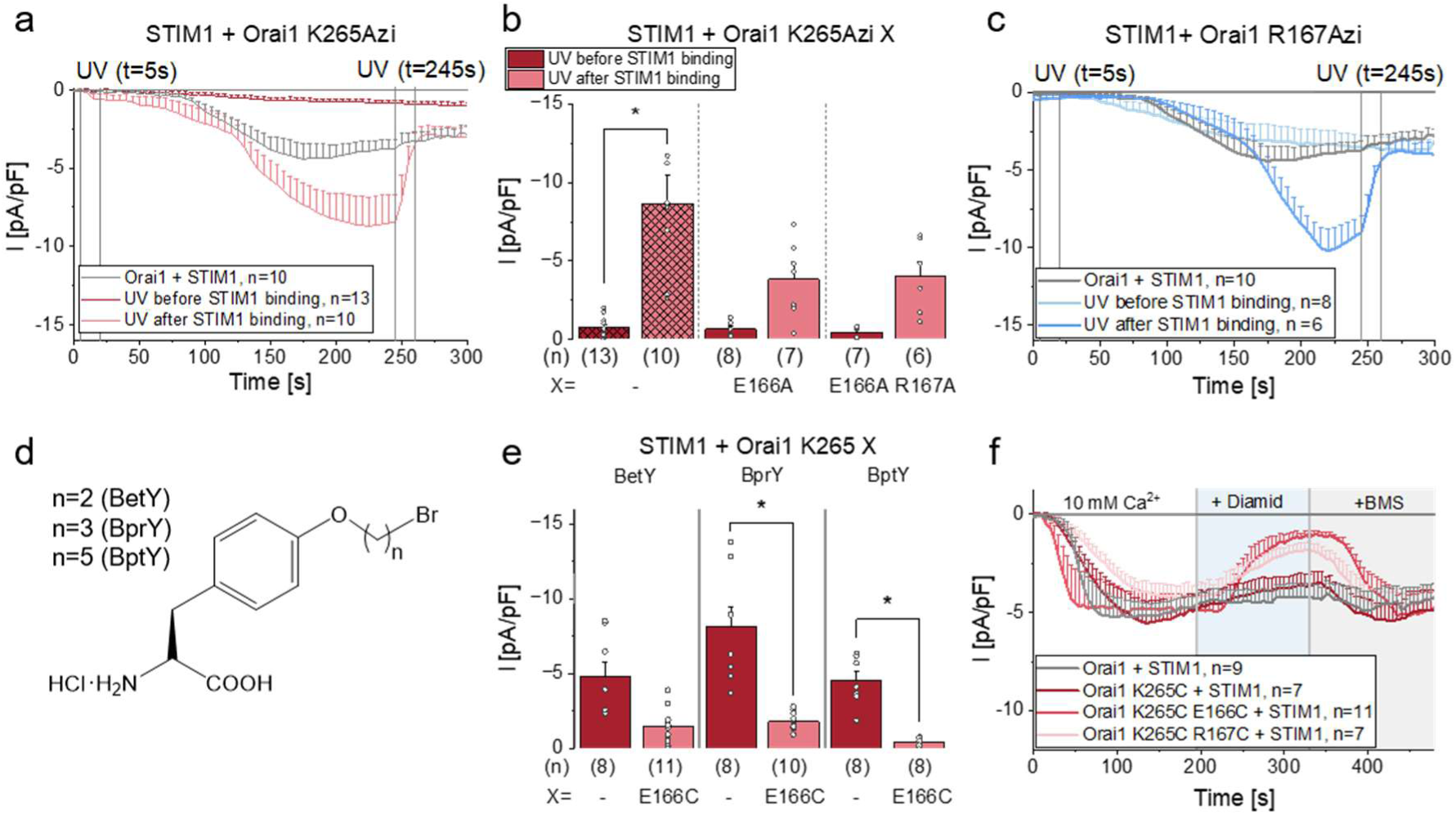
Restraining the nexus-TM3 interface via photo- and chemical crosslinking impairs STIM1-mediated maximal activation a) **c)** Time course of current densities after whole cell break-in of Orai1 WT compared to a) Orai1 K265Azi and c) Orai1 R167Azi, with 15s UV-illumination before (at t= 5s) and after (at t= 245s) STIM1-binding. **b)** Bar diagram showing maximal STIM1-mediated current densities of Orai1 K265Azi and Orai1 K265Azi E166A/E166A R167A, with 15s UV-illumination before (at t= 5s) and after (at t= 245s) STIM1-binding. Statistical significance between both conditions of UV-illumination are indicated by asterisk **\*** (p< 0.05). **d)** Chemical structure of chemical cross-linking UAAs with different linker length n. **e)** Bar diagram depicting maximal STIM1-mediated current densities of Orai1 K265-BetY/BprY/BptY and Orai1 K265-BetY/BprY/BptY E166C in the presence of STIM1. Statistical significance between single and double mutant containing a cysteine are indicated by asterisk **\*** (p< 0.05). **f)** Time course of current densities after whole cell break-in of Orai1 WT, Orai1 K265C and Orai1 K265C E166C/ R167C in the presence of STIM1. The application of diamide is indicated by the blue background (at t= 195s) and of BMS by the grey background (at t= 330s). The respective application was performed after maximal or minimal currents were reached, respectively. Single values are indicated as grey circles. Data represent mean values ± SEM of indicated number (n) of experiments. Detailed statistic values are shown in Supplementary Table 1.

Also for Orai1 R167Azi, UV-illumination before STIM1-binding led to much lower currents than STIM1-binding followed by UV-illumination, still reaching WT-like current levels (Fig. 5c). FRET-levels for R167Azi were similar to Orai1 WT regardless of the UV-exposure (Supp. Fig. 4b).

Next, we used chemical crosslinking UAAs, namely a series of tyrosine analogues containing the haloalkane bromine linked with aliphatic chains of varying length (BCnY) (Fig. 5d). We denote our UAAs BetY, BprY and BptY for a modified tyrosine with added bromoethane, bromopropane and bromopentane, respectively. The chemical crosslinking UAA binds spontaneously, covalently and selectively via a nucleophilic reaction to the thiolate of a cysteine in close proximity^29, 39, 40^ and allows us to restrain the distance between two positions (e.g. E166C K265-BCnY). Whereas Orai1 K265-BetY alone showed WT-like current levels after STIM1-activation, the STIM1-mediated current for the double mutant Orai1 K265-BetY E166C was significantly reduced (Fig. 5e, Supp. Fig. 4c). Similar results were detected for the one CH-bond-longer BprY. The double mutant Orai1 K265-BprY E166C exhibited strongly reduced currents whereas the single mutant Orai1 K265-BprY reached current levels higher than WT. Additionally, we examined the effect of BptY, three aliphatic repeat units longer than BetY, at K265 in Orai1 E166C and discovered again lower current than when integrated in Orai1 WT (Fig. 5e). Overall, all chemical crosslinking UAAs independent of side-chain length allow crosslinking with opposite cysteine, while distinct store-operated current levels in the absence of E166C might be assigned to distinct expression levels. Previous literature suggests that these chemical crosslinking UAAs can reach their counterpart within the range of up to 15Å^41^, which matches with mean distance of backbone (Cα) to backbone of 12Å ± 0.37 and with interatomic mean distance of 18Å ± 1.69 between the negatively charged carboxyl oxygen (COO^-^) of E166 and the amino nitrogen (NH ^+^) of K265 estimated from our MD simulation on the Orai1 homology model based on dOrai published in Najjar et al.^25^ (Supp. Fig. 5a)

In contrast, the double mutant Orai1 K265-BetY R167C showed a current size comparable to that of Orai1 K265-BetY alone (Supp. Fig. 4d). A reason could be that the chemical crosslinker is too short to reach R167C and therefore the nexus-TM3 interface is still able to widen/rearrange during STIM1-binding. Indeed, the longer BptY at K265 led to lower currents in combination with R167C compared to the Orai1 K265-BptY mutation alone.

To further prove that conformational rearrangements along the nexus-TM3 interface are essential for STIM1-mediated pore opening, we performed cysteine-crosslinking experiments allowing a reversible formation of a disulfide bond between two cysteines in close proximity via the addition of diamide for cysteine crosslinking and BMS (Bis(2-mercaptoethyl)sulfone) for reducing disulfide bonds back to the free sulfhydryl groups, respectively. For all four tested mutants (two controls: Orai1 and Orai1 K265C and two test mutants: Orai1 K265C E166C and Orai1 K265C R167C), STIM1-binding resulted in WT-like Ca^2+^ currents of the single and double cysteine mutants. The subsequent addition of diamide decreased current levels significantly for both Orai1 K265C E166C and Orai1 K265C R167C, likely due to formation of a cysteine crosslink, whereas no current decrease was observed for both control conditions (Fig. 5f, Supp. Fig. 4e). The break of the disulfide bond via BMS restored initial current levels for K265C E166C and K265C R167C but did not drastically affect WT and K265C (Fig. 5f). Hence, we were able to reversibly modulate STIM1-mediated currents by reversible formation of a cysteine crosslink between K265 in the nexus and an opposite residue in TM3 (E166, R167).

Overall, these results showed that crosslinking within the nexus-TM3 interface can hinder proper STIM1-mediated pore opening, likely due to restricted STIM1-mediated conformational changes involving a widening.

### STIM1-coupling to and STIM1-mediated activation of Orai1 require charges in the nexus-TM3 interface

Considering that Orai1 K265Azi E166A led to reduced STIM1-mediated coupling and activation, as well as that most residues in and opposite the nexus are charged, we pursued with analyzing the relevance of these charges in more detail (Fig. 6a). To back up our experimental protocol on charge substitutions as described below, we first investigated the apparent pKa values of the relevant residues E166, R167, H264 and K265 theoretically using both the PROPKA 3 package^42, 43^ as well as a Monte-Carlo based titration procedure based on the linearized Poisson-Boltzmann equation (MC-PBE)^44–47^ on the homology model of hOrai1^48^ based on the closed X-ray structure of dOrai (PDB: 4HKR)^14^. Orai1 E166, R167 and K265 showed monomer-independent results with apparent pKa values of 3.58, 12.94 and 10.07 in PROPKA, respectively (Supp. Fig. 6). The MC-PBE analysis revealed a clear pH independent preference of the respective charged side chains and the absence of a Henderson-Hasselbalch transition in these residues. Also, for H264, the results were monomer-independent in PROPKA, which predicted an apparent pKa of 7.14. However, in the MC-PBE approach the pKa of H264 was monomer-dependent and a strong tendency to retain the positive net charge in the imidazole ring at pH 7 with protonation probabilities ranging from 70% to 100% was reported (Supp. Fig. 6). This finding reflects considerable change with respect to the typical pKa value of 6.6 of histidine in folded proteins ^49^. Hence, our theoretical analysis shows that all relevant residues in the nexus region have a distinct preference for engaging their charged states at pH 7 and motivates us to proceed with our following mutation strategy focusing on charge deletions and charge swaps to underscore the relevance of intra- and intermolecular salt bridges in the nexus region during STIM1-Orai1 association.

**Figure 6:**
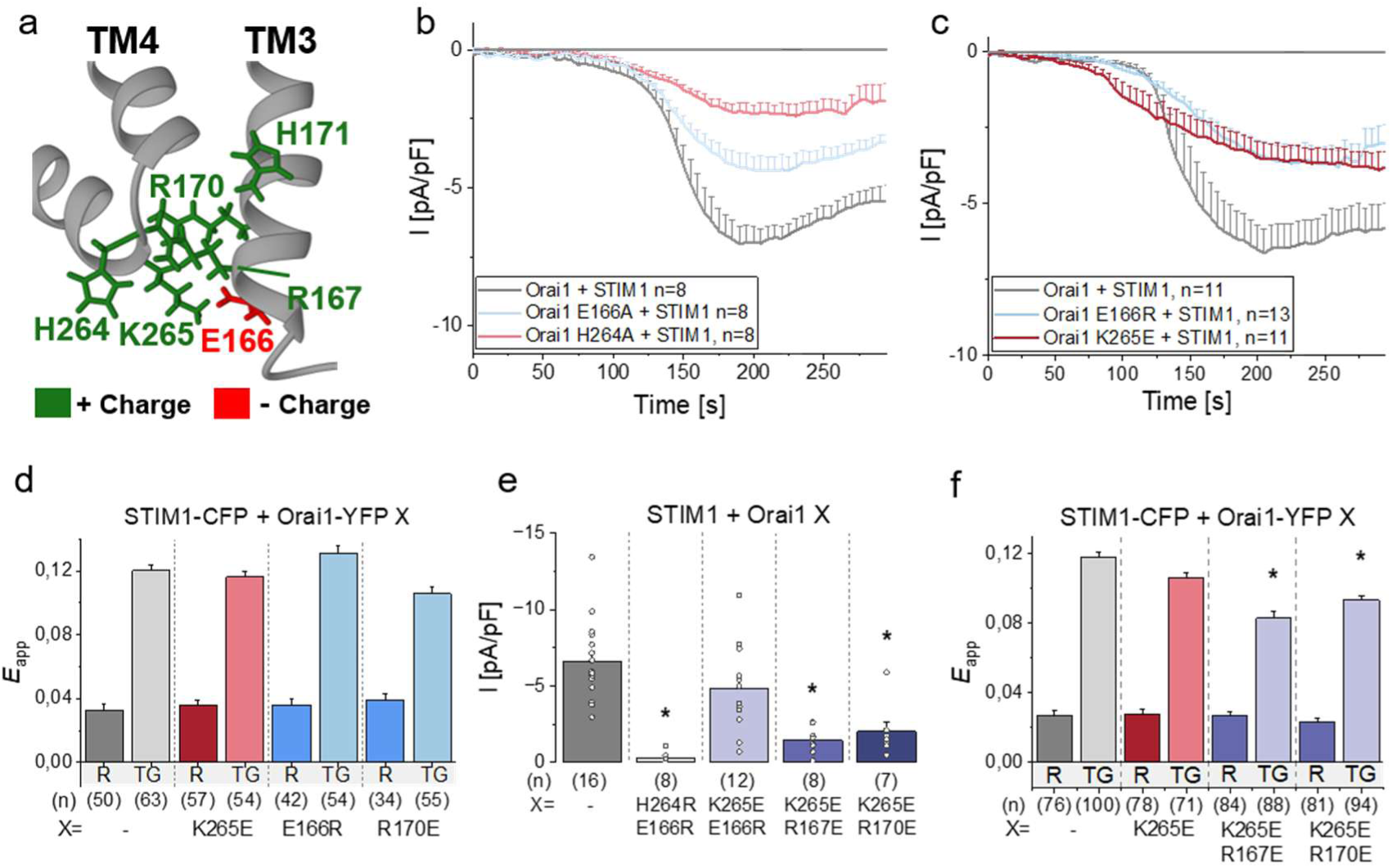
STIM1-coupling to and STIM1-mediated activation of Orai1 require charges in the nexus-TM3 interface. **a)** Representative scheme of positively (green) and negatively (red) charged residues in the Orai1 nexus-TM3 interface. **b) c)** Time course of current densities after whole cell break-in of Orai1 WT and b) Orai1 E166A/ H264A and c) Orai1 E166R/ K265E in the presence of STIM1. d) **f)** Bar diagrams depicting FRET efficiency (Eapp) detecting the interaction of STIM1-CFP with Orai1-YFP WT and d) Orai1-YFP K265E/E166R/R170E and f) Orai1-YFP K265E/ K265E R167E/R170E at resting conditions (R) and after store-depletion with 1 µM Thapsigargin (TG). Statistical significance of Orai1 mutants compared to Orai1 WT at similar conditions are indicated by asterisk **\*** (p< 0.05). **e)** Bar diagrams showing maximal current densities of Orai1 WT and Orai1 H264R E166R/K265E E166R/K265E R167E/K265E R170E in the presence of STIM1. Statistical significance of Orai1 mutants compared to Orai1 WT are indicated by asterisk **\*** (p< 0.05). Single values are indicated as grey circles. Data represent mean values ± SEM of indicated number (n) of experiments. Detailed statistic values are shown in Supplementary Table 1.

In a first step, we individually mutated charged residues in TM3 and the nexus to an alanine. Most alanine-mutants (K265, R167, R170, H171A) showed STIM1-mediated currents rather similar to Orai1 WT, except Orai1 E166A and Orai1 H264A (Fig. 6b, Supp. Fig. 7a), which exhibited significantly decreased STIM1-mediated currents. This indicates an important function in STIM1-mediated Orai1 activation of the negatively charged glutamic acid at position 166 and the positively charged histidine at position 264 (Supp. Fig. 6). Indeed, combination of H264A with neutralizing mutations in the nexus and TM3 (H264A K265A, R167A H264A) which alone retained current unaffected, reduced STIM1-mediated currents (Supp. Fig. 7b). Interestingly, double and triple mutant combinations with E166A (E166A R167A, E166A R167A K265A, E166A R167A H264A K265A) exhibited restored STIM1-mediated current levels like Orai1 WT. Other combinations (R167A K265A, R170A H171A) showed comparable or slightly enhanced STIM1-induced currents compared to WT (Supp. Fig. 7b). Nevertheless, FRET levels of Orai1-YFP E166A/ R167A K265A and E166A R167A K265A in combination with STIM1-CFP showed comparable results to STIM-CFP + Orai1 WT-YFP (Supp. Fig. 7c).

Next, we substituted a positively charged amino acid by a negatively charged one and vice versa. Here, we were indeed able to show reduced current size for Orai1 E166R as well as for Orai1 K265E, but not for Orai1 R167E and R170E (Fig. 6c; Supp. Fig. 7d). FRET-levels for these mutants were similar to WT, suggesting unaltered STIM1-Orai1 binding (Fig. 6d).

Finally, we combined charge replacement mutations in the nexus and opposite in TM3. We observed significantly lower STIM1-mediated currents for Orai1 R167E K265E and Orai1 R170E K265E than for WT (Fig. 6e). Even in FRET-experiments, the maximal FRET-levels after TG treatment were significantly reduced for the double mutants Orai1 R167E K265E and Orai1 R170E K265E (Fig. 6f). For another mutant, Orai1 E166R H264R, STIM1-mediated currents were even abolished (Fig. 6e).

In summary, charged residues at the nexus-TM3 interface are crucial for STIM1-binding and pore opening, as charged substitutions at certain positions led to decreased Orai1-STIM1-binding efficiency as well as low STIM1-mediated currents.

### Photocrosslinking UAAs located on the opposite side of the upper hydrophobic nexus exhibit strong UV-mediated current activation

In contrast to all UAA-containing mutants in the lower nexus-TM3 interface showing UV-mediated reduction of STIM1-triggered maximal currents, two mutations in the upper nexus-TM3 interface, namely H171Azi and L174Bpa depicted a strong UV-mediated current activation as revealed by our screen (Fig. 1i, Supp. Fig. 1d). Interestingly, while H171Azi is inactive without co-expression of STIM1, Orai1 L174Bpa shows UV-induced activation even in the absence of STIM1, as we previously demonstrated in Maltan et al.^26^ (Supp. Fig. 1e,f). L174 and the opposite residue L261 compose an already known hydrophobic contact essential in STIM1-binding, pore hydration and channel gating ^9, 23^. Furthermore, also position H171 has been identified as crucial for channel activation, contributing to gating by stabilizing TM3-TM4-contacts ^50^.

To obtain a better understanding of the molecular mechanisms around these residues culminating in a reversed UV-effect, namely UV-induced activation, in contrast to the observed UV-induced current decrease of lower nexus-TM3 UAA-mutants, we mutated residue L261, that is pointing towards TM3, to amino acids of different properties (Fig. 7a).

**Figure 7:**
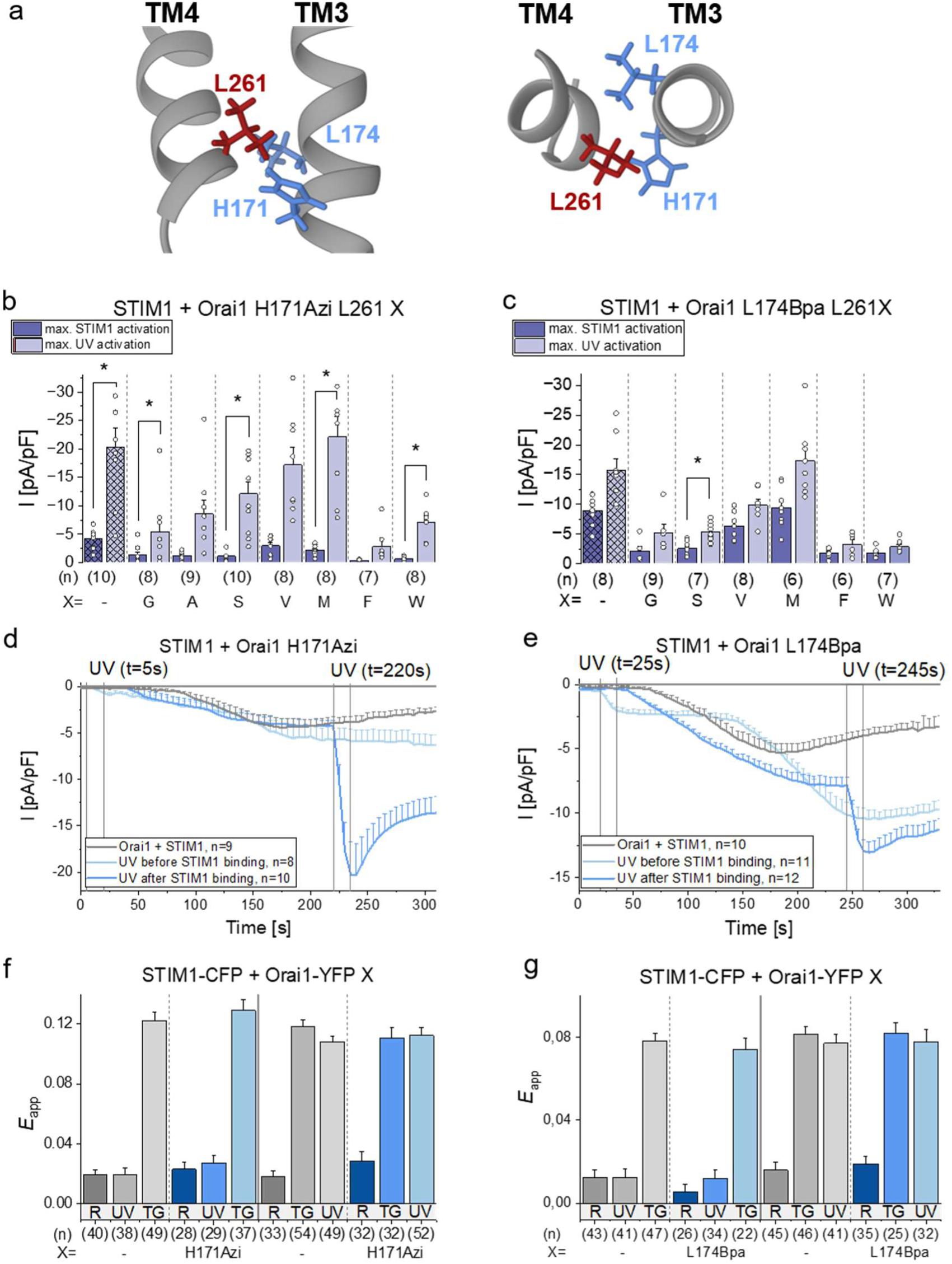
Photocrosslinking UAAs located on the opposite side of the upper hydrophobic nexus exhibit strong UV-mediated current activation. **a)** Schematic representation of the side view (left) and top view (right) of the upper nexus-TM3 interface highlighting key residues H171, L174 (TM3) and L261 (TM4). **b) c)** Bar diagrams showing maximal current densities after STIM1-activation and 15s UV-illumination of b) Orai1 H171Azi L261G/A/S/V/M/F/W and c) Orai1 L174Bpa L261G/S/V/M/F/W in the presence of STIM1. Statistical significance before and after UV-illumination are indicated by asterisk **\*** (p< 0.05). **d) e)** Time course of current densities after whole cell break-in of Orai1 WT and d) Orai1 H171Azi and e) Orai1 L174Bpa with 15s UV-illumination before (at t= 5s (d)/ 25s (e)) and after (at t= 220s (d)/ 245s (e)) STIM1-binding. **f) g)** Bar diagram showing FRET efficiency (Eapp) detecting the binding interaction of STIM1-CFP with Orai1-YFP WT and f) Orai1-YFP H171Azi and g) Orai1-YFP L174Bpa at resting conditions (R) and with 60s UV-illumination (UV) before (left) and after (right) store-depletion with 1 µM Thapsigargin (TG) triggering STIM1-binding. Data represent mean values ± SEM of indicated number (n) of experiments. Detailed statistic values are shown in Supplementary Table 1.

Mutations to glycine, alanine or serine but also to phenylalanine and tryptophan nearly abolished STIM1-mediated currents for both Orai1 H171Azi L261G/A/S/F/W and Orai1 L174Bpa L261G/S/F/W. Interestingly, for Orai1 H171Azi L261G/A/S/W and Orai1 L174Bpa L261G/S UV-illumination led to a current increase reaching WT-like or even higher levels, indicating that photocrosslinking restores the communication between the two transmembrane domains, respectively (Fig. 7b,c). As exemplarily shown for Orai1 H171Azi L261S, the UV-induced current increase only occurs in the presence and after activation of STIM1, thus, suggesting a conformational rearrangement of the upper nexus-TM3 interface after STIM1-binding (Supp. Fig. 8a).

For Orai1 H171Azi L261F and Orai1 L174Bpa L261F/W, UV-exposure was unable to significantly elevate current levels in contrast to Orai1 H171Azi/L174Bpa alone (Fig. 7b,c). Hence, the different size and/or hydrophobicity of L261-mutants seem to interfere with both, STIM1- and UV-induced activation and thus, with the communication within the upper nexus-TM3 interface.

In contrast, substitutions with either valine or methionine, both rather similar to leucine regarding hydrophobicity and size, showed STIM1-mediated current levels nearly comparable to H171Azi/L174Bpa, suggesting that the hydrophobic connection between H171Azi/L174Bpa L261V/M is still intact. Furthermore, a high UV-mediated current increase, to a rather similar level as for Orai1 H171Azi/L174Bpa, was observed, probably stabilizing this interaction by a photocrosslink (Fig. 7b,c).

Notably, the UV-mediated current size increased stepwise with increasing amino acid size from the smallest glycine to larger methionine, most similar to leucine regarding size, hydrophobicity and current level. A further increase in size like for L261F/W reverses this trend showing again smaller UV-mediated currents. Thus, the size but also the hydrophobic character of L261 seems to be critical for transferring the gating signal from the upper nexus towards the center of the channel complex (Fig. 7b,c). In accord with the critical role of hydrophobicity in the upper nexus-TM3 interface in pore opening, also STIM1-mediated currents of Orai1 L174F L261F were nearly abolished (Sup. Fig. 8b).

In addition, we investigated the effect of UV-induced photocrosslinking before STIM1-binding. Here, we observed for both Orai1 H171Azi and Orai1 L174Bpa that UV-illumination directly at the beginning of the whole cell-measurement resulted in a slight UV-mediated current increase with subsequent STIM1-mediated current activation. Orai1 H171Azi reached only WT-like levels, in contrast to when UV-illumination occurred after STIM1-binding (Fig. 7d). This suggests that, in particular around H171, conformational changes within the upper nexus-TM3 interface happen after STIM1-binding, which allow enhanced UV-induced activation of Orai1 H171Azi. Orai1 L174Bpa showed only slightly but not significantly reduced current levels when UV-light is applied before STIM1-binding compared to the reverse order of the stimuli (Fig. 7e). Hence, the covalent bond formed between L174Bpa and its potential counterpart in TM4 does not interfere with maximal STIM1-mediated activation. STIM1-binding to Orai1 H171Azi and Orai1 L174Bpa remained unaffected of the order of store-depletion and UV-induced crosslinking in FRET experiments (Fig. 7f,g).

To summarize, a hydrophobic gating interaction in the upper nexus-TM3 interface (H171/L174/L261) is essential for proper STIM1-mediated currents, as mutations of amino acids with different hydrophobicity and/or size diminish pore opening. UV-mediated photocross-linking can restore this interaction leading to strongly increased currents.

## Discussion

In this study, we report that the lower nexus-TM3 interface, which connects the Orai1 C-terminus, the major STIM1-binding site^14^, to the channel subunit, undergoes a conformational change involving its widening and contributes to proper STIM1-coupling allowing pore opening. These dynamics and interactions likely affect downstream key sites, such as the upper nexus-TM3 as well as the TM3/4-interface, contributing to the transmission of the STIM1-induced activation signal from the C-terminus^7, 13^ to the inner core of the channel, the pore. Here, our multifaceted experimental approach, combining conventional site-directed mutagenesis, incorporation of photocrosslinking UAAs, chemically crosslinking UAAs, and reversible cysteine crosslinking experiments, allowed us to gain robust insights into critical interaction partners and patterns as well as steric requirements in the nexus-TM3 interface that trigger STIM1-mediated pore opening ^17, 44^.

The incorporation of UAAs enables us to overcome limitations of conventional mutagenesis by transferring novel physicochemical properties to individual amino acids. Photocrosslinking UAAs offer the unique opportunity to study the dynamics of structure-function relationships and site-specific conformational rearrangements, as demonstrated by our previous studies ^26^. Yet, Azi exhibits lower photochemical specificity and stability than Bpa, which can lead to unspecific reactions such as ring expansion upon UV-illumination, potentially introducing steric effects, or interactions with membrane lipids ^51–53^. To circumvent this issue, we further implemented chemical crosslinking UAAs, enabling to map crosslinking partners^29^. As our experimental data showed robust and reproducible UV-induced effects, which occurred in the presence, but not or distinctly in the absence of STIM1, and are further in line with chemical and cysteine cross-linking experiments, we suggest that non-specific UV-induced crosslinking is unlikely to account for the observed functional outcomes. Notably, the most prominent effects are observed with Azi rather than Bpa, likely because Azi’s smaller size allows it to better adapt to the surroundings within the nexus-TM3 interface.

The results of our different mutagenesis approaches all consistently indicate that a widening of the nexus-TM3 interface leads to pore opening. On the one hand, this is supported by the higher STIM1-induced current densities after incorporation of a bulky amino acid, either a photocrosslinking UAA or an aromatic amino acid, in particular at K265. On the other hand, cross-linking (photo-, chemical, cysteine crosslinking) at certain positions at the nexus-TM3 interface leads to a reduction in STIM1-mediated current densities. While alanine-substitutions only at some sites reduce store-operated current activation, either due to narrowing of the nexus-TM3 interface and/or reduced STIM1-coupling, UV-light triggers current reduction to WT- or even smaller levels. Overall, this indicates that crosslinking decreases the nexus-TM3 distance, and our crosslinking assays, in particular chemical and cysteine crosslinking, enabled us to map a contact area contributing to the maintenance of the closed state. In turn, these findings suggest that STIM1 can cause a widening of this interface to trigger signaling to pore opening. In support, GoF-mutants containing the UAA mutation at critical sites in the nexus-TM3 interface (e.g. K265) do not respond to UV-light, likely due to a GoF-mutation-induced conformational change, such as widening, at the nexus-TM3 interface that distances potential crosslinking partners from each other. Our observed slight tendency for UV-induced current reduction in the presence of STIM1 (e.g. of Orai1 K265Azi V181K) suggests some STIM1-induced structural rearrangements, although the open conformation is already set. This supports previous evidence that STIM1-binding fine-tunes pore opening ^38^. Under physiological conditions, the widening of the nexus-TM3 interface could be caused by the bending of the nexus node (aa 261-265) after STIM1-binding. These findings are in line with previous reports showing that STIM1-binding and/or pore opening are linked to conformational changes of the C-terminus ^9, 14, 15^. Furthermore, our previous studies demonstrate that the TM3-TM4 interface in particular around the V181 position and directly connected to the nexus-TM3 interface undergoes widening to induce pore opening ^25, 37^. Hence, it is tempting to speculate that the signal of STIM1-binding is transmitted via the nexus-TM3- to the TM4-TM3 interface, which triggers further transmission to the center of channel complex.

Beyond the widening of the nexus-TM3 interface, additional conformational rearrangements may still occur. In contrast to Orai1 R167Azi, Orai1 H171Azi, and Orai1 K265Azi, which exhibit reduced maximal currents when UV-light is applied prior to STIM1-coupling, Orai1 L174Bpa demonstrates nearly comparable currents regardless of the order in which these stimuli are applied. The fact that the putative UV-induced crosslink formation of L174Bpa with TM4 still permits high current activation suggests that STIM1, in addition to inducing a widening of the nexus-TM3 interface, may also trigger other structural rearrangements.

While the incorporation of the photocrosslinking UAA at K265 and R167 or an aromatic amino acid at K265 in the nexus resulted in significantly increased currents, store-operated currents of other canonical substitutions of different properties at R167 showed no side-chain dependence. A possible reason could be that R167 is not directly pointing towards the nexus but rather pointing slightly out of the nexus-TM3 interface as revealed by investigations of the hOrai1 homology model (Fig. 3a) as well as our hOrai1 MD simulations^25^ (Supp. Fig. 5b). Hence, changes of the amino acid size of position 167 does not directly influence the distance in the respective interface.

Several of our results hint on a potential role of the nexus-TM3 interface in STIM1-coupling. While several aromatic substitutions enhanced STIM1-mediated Orai1 currents, others (e.g. K265W, R167F K265F) did not strongly or not at all enhance STIM1-induced currents. FRET experiments showed that R167Y K265Y exhibits reduced STIM1-coupling. This result is in accord with our previous report, in which we demonstrated impaired STIM1-coupling for some constitutively active Orai1 mutations located at the TM3-TM4 interface ^37^ (Fröhlich et al. unpublished). Heterogenous effects of various A-substitutions in UAA-containing Orai1 mutants cannot only be attributed to changes along the nexus-TM3 interface but could potentially also affect STIM1-coupling. Indeed, several A-substitutions (E166A, E166A R167A, E166A R170A) in Orai1 K265Azi reduced STIM1-coupling. Furthermore, our findings point to a critical impact of charges within this interface in the STIM1-mediated activation. The insertion of reversed charges (E166, K265) or alanine (E166) resulted in reduced STIM1-mediated currents for multiple single or double mutants probably due to strengthened or weakened electrostatic interactions. For Orai1 K265E R167E/R170E even STIM1-Orai1 binding was negatively affected. Hence, charges within the nexus-TM3 interface play a critical role in fine-tuning STIM1-coupling and pore opening. It remains unclear whether these charges in the nexus-TM3 interface affect STIM1-binding directly or indirectly. As cysteine crosslinking of E166C and K265C works in the STIM1-bound state, we suggest a more indirect effect of the charges on STIM1-binding.

Taking a closer look on the structure reveals that at least in three subunits K265 and E166 point to each other (Supp. Fig. 5a,9). Thus, crosslinking of at least three subunits leads to robust current reduction. However, it is tempting to speculate that STIM1-binding triggers a reorientation leading to more E166-K265 pairs pointing to each other, thus making up robust crosslinking induced functional effects. However, this requires further proof by a structural resolution of STIM1 and Orai1 in complex. In addition to cysteine crosslinking induced current reduction of Orai1 E166C K265C, we previously demonstrated that also Orai1 K78C E166C shows diamide-induced current reduction ^54^. The reason for this could be that at least in three subunits both, E166 and K265 point also to K78 (Supp. Fig. 5a). Moreover, these findings indicate that changes in charges in this area can affect the local surroundings and consequently impact STIM1-coupling and -mediated activation. Moreover, Orai1 E166C could be activated by crosslinking with STIM1 L402C^55^, supporting the critical role of E166 and its environment in STIM1-coupling.

Our primary screen of the nexus-TM3 interface revealed two UAA-mutants in the upper nexus (H171Azi, L174Bpa) showing STIM1-mediated activation followed by strong UV-induced current activation (Fig. 1i, Supp. Fig. 1d). Both STIM1-mediated and UV-induced activation depend on the amino acid properties at position 261. While small or aromatic amino acids allow no or only weak store-operated activation, amino acids (V, M) with similar size/hydrophobicity like leucine show WT-like STIM1-mediated current levels. This indicates that hydrophobic interactions in the upper nexus-TM3 interface are crucial for maintaining the closed state but also for STIM1-induced activation. In line with the role of hydrophobicity in the upper nexus, two phenylalanines at L174 and L261 abolish STIM1-mediated activation. In accord with our results, other contact pairs close to the nexus-TM3 interface (F178-F257^17^, A175-Y258^50^) have been reported to play a critical role in the pore opening mechanism.

UV-induced currents of both Orai1 H171Azi and Orai1 L174Bpa correlate with an increase in side-chain size at position 261 in accord with our observations that widening of the nexus-TM3 and the TM3-TM4 interfaces contributes to pore opening. Only aromatic amino acids (F, W) at 261 do not allow strong UV-induced activation, potentially due to steric hindrance or hydrophobic interactions generally impairing activation. A covalent interaction formed starting from H171 seems to stabilize the open state, which is in accord with our previous finding on the critical role of the TM3-TM4 interface in Orai1 pore opening ^25^.

Interestingly, in the absence of STIM1, Orai1 L174Bpa, but not H171Azi showed UV-induced activation. Taking a closer look at the closed state Orai1 structure reveals that L174Bpa is always opposite S260/L261 in TM4, while H171 is pointing out of the nexus-TM3 interface or to the C-terminus (Supp. Fig. 5c,9). UV-induced current enhancements after maximal STIM1-induced activation of Orai1 H171Azi points to a STIM1-binding induced structural rearrangement of C-terminus and TM4 potentially making residues in TM4 accessible for photocross-linking. This behaviour is reflected by corresponding double mutants containing L261 mutated to amino acids with different properties. Interestingly, the open state structure indicates that H171 might still be oriented outward of the nexus-TM3 interface (Supp. Fig. 9), which is in contradiction to our experimental data, and highlights the need for structural resolution of STIM1 complexed with Orai1.

In summary, we discovered that the nexus-TM3 interface plays a critical role in the STIM1-mediated pore opening process of Orai1. With the use of photocrosslinking and chemical crosslinking UAAs as well as site-directed mutagenesis, we showed that a widening of the lower nexus-TM3 interface is essential for proper channel activation by STIM1. Furthermore, the contacts formed at the upper nexus-TM3 interface are necessary for signal transduction and pore opening. Our experiments support the findings of Zhou et al.^23^ that the binding of STIM1 to the Orai1 C-terminus results in the flexion of the lower nexus, thus inducing a widening of the nexus-TM3 interface. This conformational rearrangement further enables the formation of gating interactions in the upper nexus-TM3 interface, that stabilize the opening conformation of Orai1 and transfer the opening signal towards the more central TM-domains. Along with this, the inner TM3-TM4-TM2’ interface widens, as we discovered in our previous publication^25^, thus transmitting the opening signal towards the central pore of the channel. Our studies provide a further basis for continuing studies of our previous findings^33^ on the role of isoform-specific features of the TM3-TM4 interfaces in fine-tuning Orai pore opening.

Elucidating the precise conformational dynamics during STIM1-mediated Orai1 activation remains a fundamental challenge in the field. Here, we uncover key mechanistic principles governing pore opening by combining site-specific photo- and chemical crosslinking with conventional mutagenesis. The discovered results gained by different approaches are all in line and supporting the critical role of the nexus-TM3 interface in pore opening and supplement previous important findings ^23, 25, 50^. This study again highlights how the use of unnatural amino acids is suitable to uncover novel CRAC channel dynamics. Our innovative methods facilitate the detailed discovery of conformational events happening during pore opening at amino acid level, thus opening new possibilities of future drug design and therapeutic strategies.

## Methods

### Molecular biology

For C-terminal fluorescent labeling of human Orai1 (accession number NM_032790; provided by A. Rao’s laboratory), the gene was cloned into the pEYFP-N1 expression vector (Clontech) using XhoI and BamHI restriction sites. Site-directed mutagenesis and amber stop codon (TAG) insertion for site-specific UAA-incorporation were carried out using the QuikChange™ XL kit (Stratagene), with the corresponding Orai1 construct as a template. Corresponding Primer sequences were purchased from Eurofins Genomics. Human STIM1 (accession number NM_003156), N-terminally tagged with ECFP, was kindly provided by T. Meyer (Stanford University). STIM1 point mutations were introduced using the QuikChange™ XL kit as well. All constructs were verified by DNA sequencing (Eurofins Genomics/Microsynth).

Humanized aminoacyl-tRNA synthetase/tRNA pairs for the recognition of unnatural amino acids azido-L-phenylalanine, benzoyl-L-phenylalanine and BCnY were purchased from Addgene (#105829, #155342 and #155343).

### Cell culture and transfection

Human embryonic kidney 293 -T (HEK293-T) cells (#ACC635, DSMZ) were cultured in DMEM, supplemented with L-glutamine (2mM), streptomycin (100μg/ ml), penicillin (100 units/ml) and 10% fetal calf serum at 37°C in a humidity-controlled incubator with 5% CO2.

Transient transfection of HEK293-T cells was performed using TransFectin™ Lipid Reagent (Bio-Rad; 2.5 μL per transfection). Standard plasmid ratios were 1 μg Orai1: 2 μg tRNA/aaRS; 1 μg STIM1 for co-expression. Experiments were conducted 24 h post-transfection in wild-type HEK293-T.

Throughout all experiments, wild-type or mutant Orai1-YFP constructs were used, along with STIM1-CFP where applicable. Media of transfected cells was supplemented with 1 mM photo-crosslinking unnatural amino acids (azido-L-phenylalanine or benzoyl-L-phenylalanine; BACHEM, dissolved in 0.5 M NaOH) and 0.3 mM chemical crosslinking unnatural amino acids (BCnY, synthesized by our collaboration partner M. Waser according to Xiang et al.^39^, dissolved in DMSO). Mycoplasma contamination was routinely tested using the VenorGeM Advanced Mycoplasma Detection Kit (Minerva Biolabs).

### Electrophysiology

Electrophysiological recordings were performed using an inverted microscope (Zeiss Axiovert 200), an Axopatch 200B amplifier (Molecular Devices), a PatchStar micromanipulator (Scientifica), and a Lumencor Spectra III light engine. Electrophysiological records were acquired with pClamp11 software. 2.2 mW/cm² UV-intensity was used within all experiments. Whole-cell patch-clamp recordings were conducted at room temperature (20°C) using an Ag/AgCl reference electrode. Cells, grown in petri dishes were reseeded onto poly-L-lysine-coated coverslips 4 h to 18 h post-transfection in medium containing 0.1 mM Ca²⁺ and used for measurements 6–10 h later. For time-course and I/V measurements, voltage ramps from –90 mV to +90 mV over 1 s were applied every 5 s starting from a holding potential of 0 mV. Fast Ca²⁺- dependent inactivation (FCDI) was assessed using voltage steps to –70, –90, or –110 mV for 2000 ms starting from a holding potential of 0 mV.

Passive store-depletion was triggered via the internal pipette solution containing (in mM): 145 Cs-methane sulfonate, 20 EGTA, 10 HEPES, 8 NaCl, 3.5 MgCl₂, pH 7.2. The standard extra-cellular solution contained (in mM): 145 NaCl, 10 HEPES, 10 CaCl₂, 10 glucose, 5 CsCl, 1 MgCl₂, pH 7.4. For cysteine crosslinking experiments 1 mL of a 1 mM Diamide in 10 mM Ca^2+^ solution was used for disulfide bond formation and 1 mL of a 5 mM BMS (Bis(2- mercaptoethyl)sulfone) in 10 mM Ca^2+^ solution was used for bond-cleavage. Solution exchange during the experiment was performed using a Thomas Wisa perfusion pump.

Applied voltages were not corrected for the liquid junction potential, which was determined as +12mV. Leak currents were subtracted either from initial baseline traces after whole-cell break-in or from 10 µM La³⁺-blocked traces at the end of the experiment.

### Confocal FRET Fluorescence Microscopy

Confocal FRET imaging was performed at room temperature (20°C) 18–24 h after transfection. The standard extracellular solution containing (in mM): 145 NaCl, 5 KCl, 10 HEPES, 10 glucose, 1 MgCl₂, 2 CaCl₂, pH 7.4. For Ca²⁺ store-depletion, a Ca²⁺-free extracellular solution containing 1 μM Thapsigargin was used. Imaging was conducted using a CSU-X1 real-time confocal system (Yokogawa Electric Corporation, Japan) equipped with two CoolSNAP HQ2 CCD cameras (Photometrics, AZ, USA), a dual-port adapter dichroic, 505lp; cyan emission filter, 470/24; yellow emission filter, 535/30; Chroma Technology Corporation, VT, USA), and two diode lasers (445 nm and 515 nm; Visitron Systems). The system was mounted on an Axio Observer.Z1 inverted microscope (Carl Zeiss) and placed on an anti-vibration table (Vision IsoStation, Newport). A Thomas Wisa perfusion pump enabled continuous extracellular solution exchange during the experiment. Image acquisition and system control were carried out via the VisiView software (v2.1.4, Visitron Systems). The illumination times for individual sets of images (CFP, YFP, FRET) that were recorded consecutively with a minimum delay were kept in a range of 100–300 ms. To account for spectral bleed-through and cross-excitation, correction factors for YFP cross-excitation (α) and CFP crosstalk (β) were determined daily using control samples expressing only CFP or YFP. FRET analysis was limited to pixels with CFP:YFP ratios between 0.1:10 and 10:0.1. After this threshold determination as well as background signal subtraction, the apparent FRET efficiency *E*_app_ was calculated on a pixel-to-pixel basis using a custom MATLAB routine (v7.11.0, MathWorks, Inc., MA, USA) and the following equation:

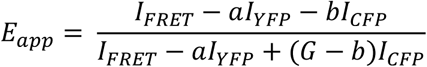

where *I_FRET_*, *I_YFP_* and *I_CFP_* denote the intensities of the FRET, YFP and CFP images, respectively. *G* denotes a microscope-specific constant parameter that was experimentally determined as 2.75 ^9, 19^.

### Molecular Modeling and structural analysis

We retrieved the all-atom homology model of hOrai1 published by Frischauf et al.^48^ from the model archive with accession code “ma-akdjp” ^56^. We defined the transmembrane region by focusing on the average z-coordinate of the R91 alpha carbon in all monomers at the cytosolic mouth of the Orai1 pore. From this point, we defined a membrane vector in z-direction with 3 nm length. This slab domain will be treated as an implicit membrane with εr= 2 throughout our MC-PBE approach ^44–46^. The protein domain was defined by its enveloping Connolly surface, which is generated by rolling a spherical water probe with a radius of 0.14 nm over the Van-der-Waals surface of hOrai1^57^. The aqueous environment is modelled with a dielectric constant εr= 80, the protein itself with εr= 4. We defined the Debye length of our system with about 0.8 nm by providing a physiological concentration of monovalent ions 0.15 M in our aqueous domain at a temperature of 300 K ^58^. In our next step, we aim at the numerical solution of the linearised Poisson-Boltzmann equation using TAPBS^46^ on a simple cubic grid for every titratable side chain in isolation and, separately, in its native protein environment. Starting from a coarse grid with 16.1 nm edge length and a grid constant of 0.3 nm, we obtained our final data from a calculation on a grid with a grid constant of 0.025 nm using the same edge length. By virtue of a thermodynamic cycle analysis^47^, we can infer the change of Gibbs free energy due to protonation of a single side chain in the protein. We use these Gibbs free energies to perform a Monte-Carlo simulation using the Metropolis-Hastings algorithm^59^ implemented in Karlsberg 2^45^ to obtain an optimal protonation pattern in the canonical ensemble of the hOrai1 channel between pH 5 and pH 9 at 300 K. These patterns are defined by microscopic protonation probabilities of single titratable side chains, which can be used to assess the preferred titration states and effective pKa values of K78, E166, K265, R167, H264 in all hOrai1 monomers at pH 7.

To control and compare our findings from the MC-PBE approach, we started an independent analysis of effective pKa values in hOrai1 with PROPKA3^42, 43^ using the code provided on the GitHub repository^60^ and pre-defined standard settings with the very same homology model structure ^48, 56^. We provide the PROPKA3 output as Supplementary File 1.

Structural analyses were performed on published molecular dynamics trajectories of the homology model of Orai1 WT based of the closed dOrai X-ray structure ^61^. Our distance measurements were carried out on random samples from the equilibrated trajectory parts. Snapshot number 122 was used for the structural analysis carried out in Supp. Fig. 5.

### Statistics and Reproducibility

Data are presented as mean ± SEM, with (n) indicating the number of independent experiments. For statistical comparison between two independent groups, the Mann–Whitney test was applied, with *p* < 0.05 considered statistically significant. Variance homogeneity was assessed using Levene’s test. If the assumption was fulfilled, one-way ANOVA test was used for the statistical comparison of multiple independent samples using the F-distribution, followed by Fisher’s least significant difference (LSD) post hoc test. If the assumption was not fulfilled, Welch’s ANOVA followed by Games–Howell post hoc test was used. Normality of the data was verified using the Shapiro–Wilk test or, alternatively, the one-sample Kolmogorov–Smirnov test. All corresponding *F* and *p* values for the Main Figures are listed in Supplementary Table 1.

All experiments were carried out at least on two different days in paired comparisons leading to similar results. Cell images were taken for each experiment showing comparable cellular distribution of the respective proteins/mutants, as shown in one representative image in the respective figures.

## Data Availability

Source data will be uploaded to zenodo upon acceptance.

## Acknowledgements

We thank S. Buchegger for excellent technical assistance. We thank C.Humer for critically proofreading the manuscript. This research was funded by the Austrian Science Fund (FWF) projects [doi:10.55776/P32851], [doi:10.55776/ P35900] and [doi:10.55776/P36202] to I.D and [doi:10.55776/PAT 6871323] to A.T. For open access purposes, the author has applied a CC BY public copyright license to any author-accepted manuscript version arising from this submission.

## Author contributions

JS and ID conceived and coordinated the study and wrote the paper. JS, HN, and TR performed and analyzed electrophysiological experiments. MP, AS carried out fluorescence microscopy experiments. JS, YN, SH, MF, DT contributed to molecular biology. VA, SW, HK performed and supervised computational studies. CM, WN and MW synthesized chemical cross-linking UAAs. All authors reviewed the results and approved the final version of the manuscript.

## Supplementary information

includes 9 Supplementary Figures (Supp. Fig. 1-9), 1 Supplementary Table and 1 Supplementary File.

Supplementary Figure 1: UV-triggered inhibition of Orai1 currents via photocrosslinking UAAs introduced in the nexus and in TM3 is dependent on the presence of STIM1.

Supplem Alanine substitutions in TM3 opposite to the nexus-UAA do not prevent UV-induced current inhibition, but affect STIM1-mediated activation.

Supplementary Figure 3: Constitutively active Orai1 mutations interfere with UV-induced current inhibition upon UAA-insertion at the nexus–TM3 interface.

Supplementary Figure 4: Photo- and chemical crosslinking at the nexus–TM3 interface restricts STIM1-mediated channel opening.

Supplementary Figure 5: Orientation of key residues in the nexus-TM3 interface is critical for UV-induced photocrosslinking.

Supplementary Figure 6: Titration states of E166, R167, H264 and K265 based on the closed state of Orai1.

Supplementary Figure 7: Charged residues along the nexus-TM3 interface are required for STIM1-mediated Orai1 channel gating.

Supplementary Figure 8: Interactions in the upper hydrophobic nexus-TM3 interface are essential for STIM1-mediated Orai1 pore opening.

Supplementary Figure 9: The channel conformation of the nexus-TM3 interface of the closed and the open state of Orai1.

Supplementary Table 1: Results of statistical tests. Supplementary File 1: Results of effective pKa values in hOrai1.

