## Supplementary Figures for "Distances and charges along the Orai1 nexus-TM3 interface control STIM1-binding and pore opening"

### Supplementary Information

#### (1) Supplementary Figures 1-9

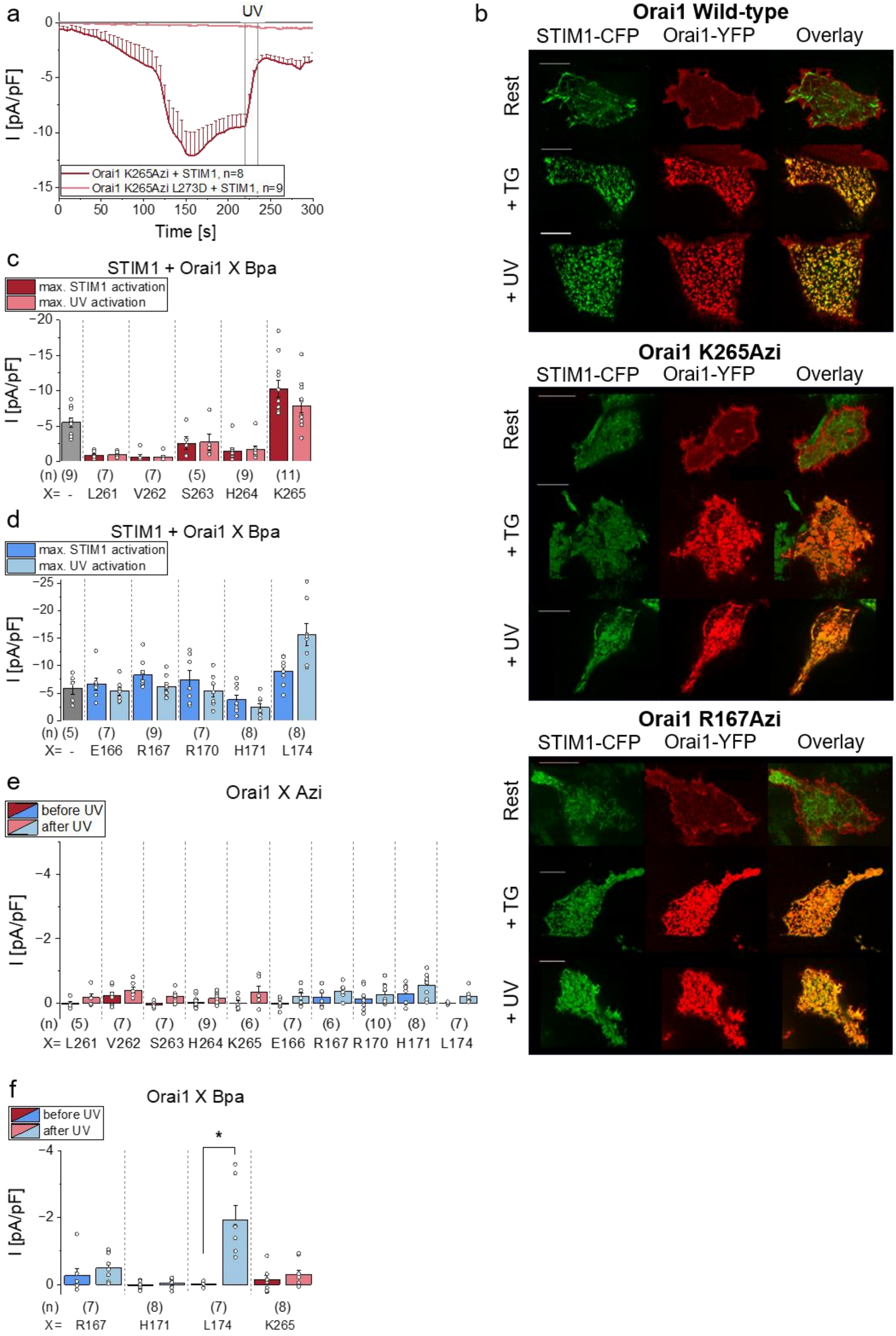
a
UV
I [pA/pF]
Time [s]
Orai1 K265Azi + STIM1, n=8
Orai1 K265Azi L273D + STIM1, n=9
b
Orai1 Wild-type
STIM1-CFP
Orai1-YFP
Overlay
Rest
+ TG
+ UV
Orai1 K265Azi
Orai1 R167Azi
c
STIM1 + Orai1 X Bpa
max. STIM1 activation
max. UV activation
(n) (9) (7) (7) (5) (9) (11)
X= - L261 V262 S263 H264 K265
d
STIM1 + Orai1 X Bpa
(n) (5) (7) (9) (7) (8) (8)
X= - E166 R167 R170 H171 L174
e
Orai1 X Azi
before UV
after UV
(n) (5) (7) (7) (9) (6) (7) (6) (10) (8) (7)
X= L261 V262 S263 H264 K265 E166 R167 R170 H171 L174
f
Orai1 X Bpa
(n) (7) (8) (7) (8)
X= R167 H171 L174 K265

**Supp. Figure 1: UV-triggered inhibition of Orai1 currents via photocrosslinking UAAs introduced in the nexus and in TM3 is dependent on the presence of STIM1**

**a)** Time course of current densities after whole cell break-in of Orai1 K265Azi and Orai1 K265Azi L273D in the presence of STIM1. UV-light is applied for 15s at t= 220s, after maximum STIM1-mediated currents were reached. **b)** Confocal fluorescence microscopy images of representative cells co-expressing STIM-CFP with Orai1-YFP WT, Orai1-YFP K265Azi and Orai1-YFP R167Azi and their overlay at resting conditions (R), after store-depletion via 1 µM Thapsigargin (TG) and after 60s UV-illumination (UV). White bars indicate 5 µm. **c) d)** Bar diagram showing maximal current densities after STIM1-activation and after 15s UV-illumination of c) Orai1 nexus-mutants and d) Orai1 TM3-mutants containing Bpa. **e) f)** Bar diagram showing maximal current densities of Orai1 nexus-TM3 mutants containing e) Azi or f) Bpa before and after 15s UV-illumination in the absence of STIM1. Statistical significance before and after UV-illumination are indicated by asterisk * ($p < 0.05$). Single values are indicated as grey circles. Data represent mean values ± SEM of indicated number (n) of experiments.

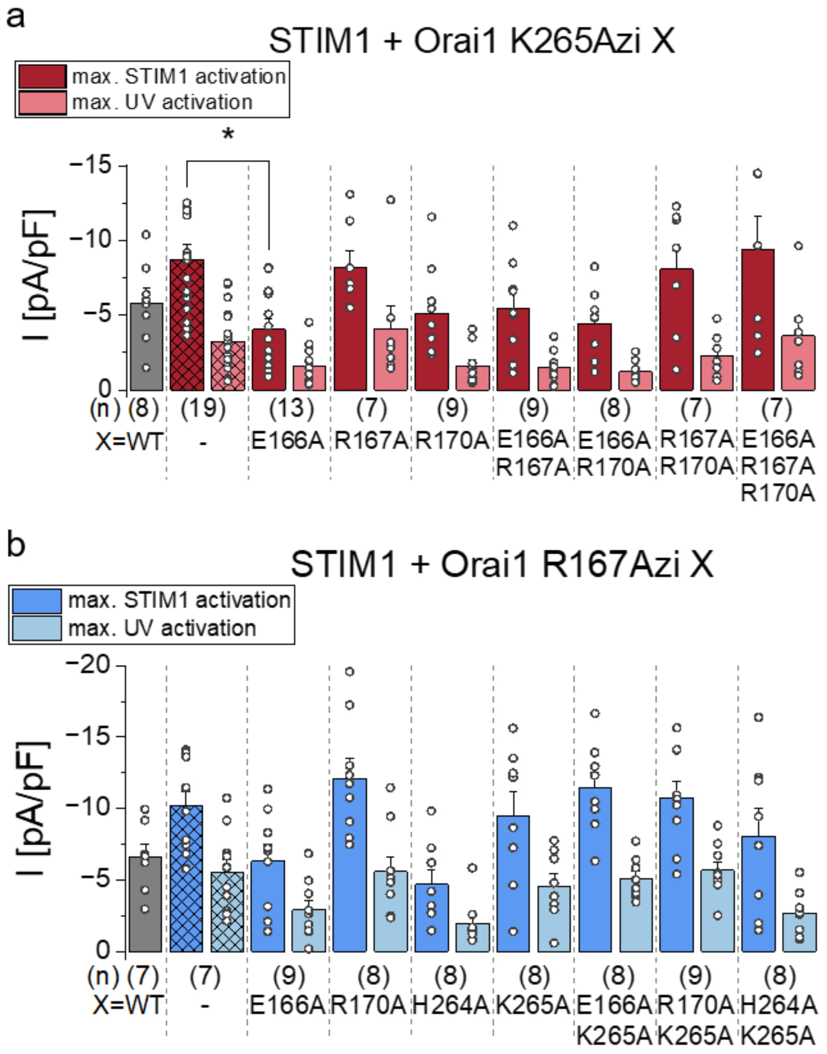


**Supp. Figure 2: Alanine substitutions in TM3 opposite to the nexus-UAA do not prevent UV-induced current inhibition, but affect STIM1-mediated activation**

**a) b)** Bar diagrams exhibiting maximal current densities after STIM1-activation and after 15s UV-illumination of a) Orai1 K265Azi-TM3-mutants and b) Orai1 R167Azi-nexus-TM3 mutants compared to STIM1-induced Orai1 WT currents. Statistical significance of Orai1 mutants compared to Orai1 K265Azi currents obtained in the presence of STIM1 at similar conditions are indicated by asterisk * ($p < 0.05$). Single values are indicated as grey circles. Data represent mean values ± SEM of indicated number (n) of experiments.

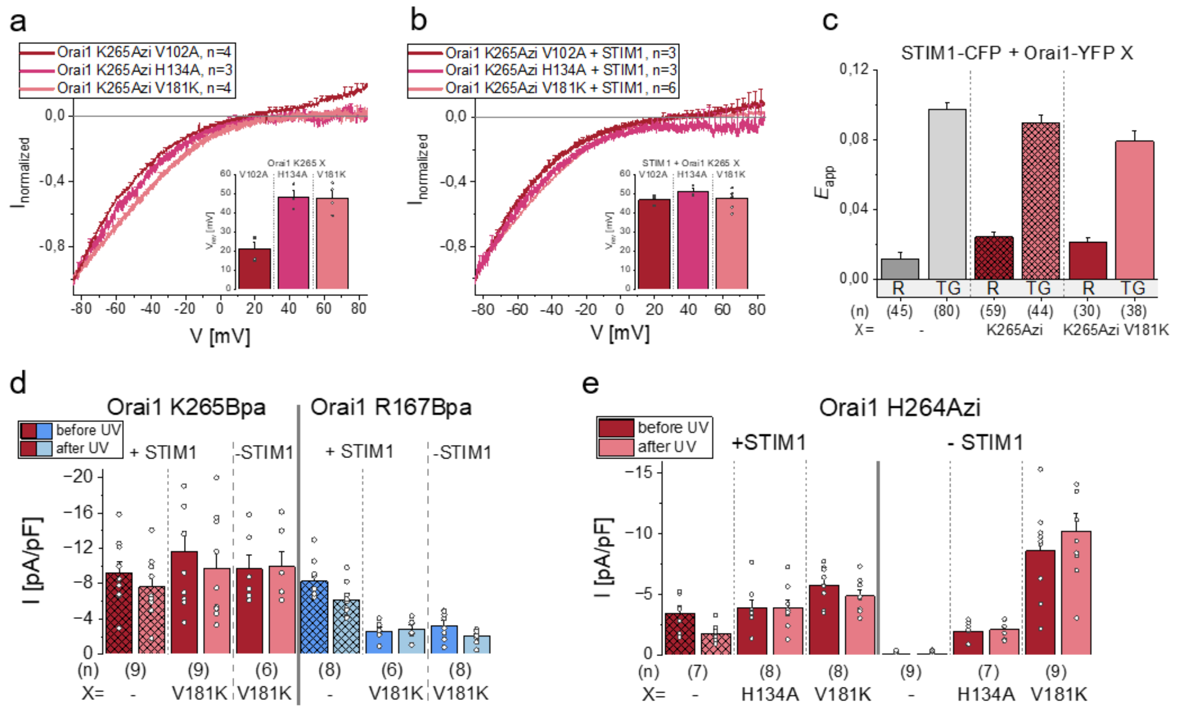


**Supp. Figure 3: Constitutively active Orai1 mutations interfere with UV-induced current inhibition upon UAA-insertion at the nexus–TM3 interface**

**a) b)** Normalized I/V relationship of Orai1 K265Azi V102A/H134A/V181K a) in the absence of STIM1 b) in the presence of STIM1. Inlet represents the reversal potential ($V_{rev}$) of a) constitutively active Orai1 currents or b) of STIM1-mediated Orai1 currents. **c)** Bar diagrams showing FRET efficiency ($E_{app}$) detecting the binding interaction of STIM1-CFP with Orai1-YFP WT and Orai1-YFP K265Azi/K265Azi V181K at resting conditions (R) and after store-depletion with 1 µM Thapsigargin (TG). **d) e)** Bar diagrams showing maximal current densities before and after 15s UV-illumination of d) Orai1 K265Bpa, Orai1 R167Bpa and Orai1 K265Bpa/R167Bpa V181K and of e) Orai1 H264Azi and Orai1 H264Azi H134A/V181K in the presence and in the absence of STIM1. Single values are indicated as grey circles. Data represent mean values ± SEM of indicated number (n) of experiments.

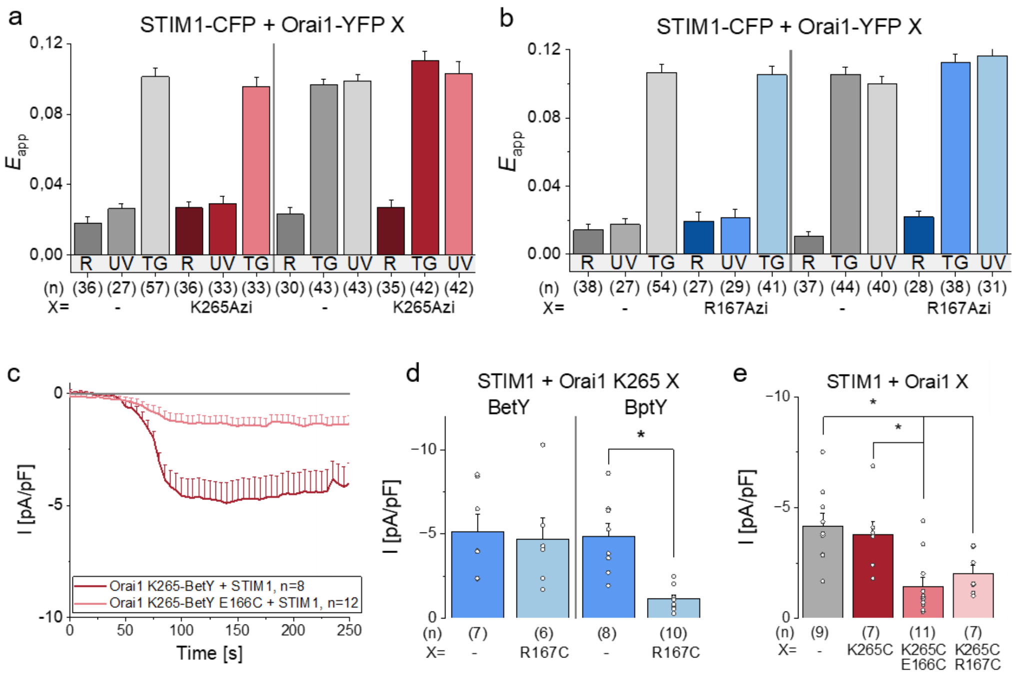


**Supp. Figure 4: Photo- and chemical crosslinking at the nexus–TM3 interface restricts STIM1-mediated channel opening**

**a) b)** Bar diagrams showing FRET efficiency ($E_{app}$) detecting the interaction of STIM1-CFP with Orai1-YFP WT and a) Orai1-YFP K265Azi and b) Orai1-YFP R167Azi at resting conditions (R) and with 60s UV-illumination (UV) before (left) versus after (right) the application of 1 µM Thapsigargin (TG) triggering store depletion and STIM1-binding. **c)** Time course of current densities after whole cell break-in of Orai1 K265-BetY and Orai1 K265-BetY E166C in the presence of STIM1. **d) e)** Bar diagram depicting maximal STIM1-mediated current densities of d) Orai1 K265Azi-BetY/BptY and Orai1 K265-BetY/BptY R167C and e) Orai1 WT, Orai1 K265C and Orai1 K265C E166C/R167C in the presence of STIM1. Statistical significance between corresponding mutants are indicated by asterisk * ($p < 0.05$). Single values are indicated as grey circles. Data represent mean values ± SEM of indicated number (n) of experiments.

a

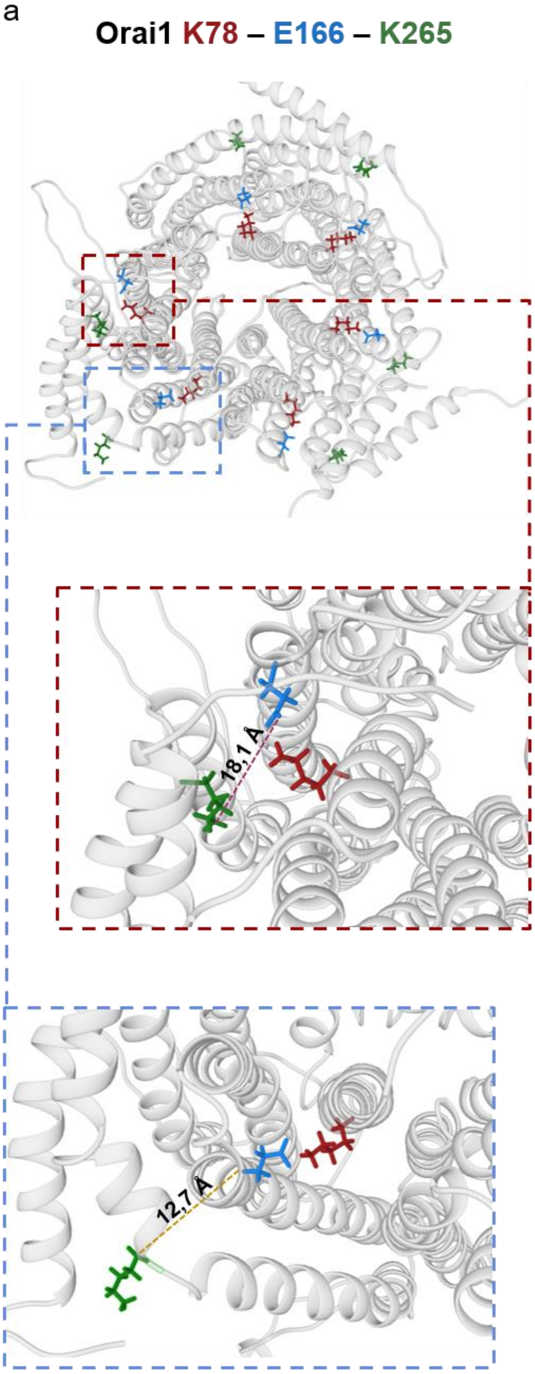


| | distance (Å) | |
|---|---|---|
| Subunit | K265 Cα - E166 Cα | K265 $N_Z$ - E166 $O_{E2}$ |
| 1 | 13,02 | 22,713 |
| 2 | 11,854 | 14,1 |
| 3 | 14,147 | 22,373 |
| 4 | 12,916 | 12,64 |
| 5 | 11,673 | 18,696 |
| 6 | 12,798 | 17,955 |
| Mean | 12,735 | 18,08 |

b

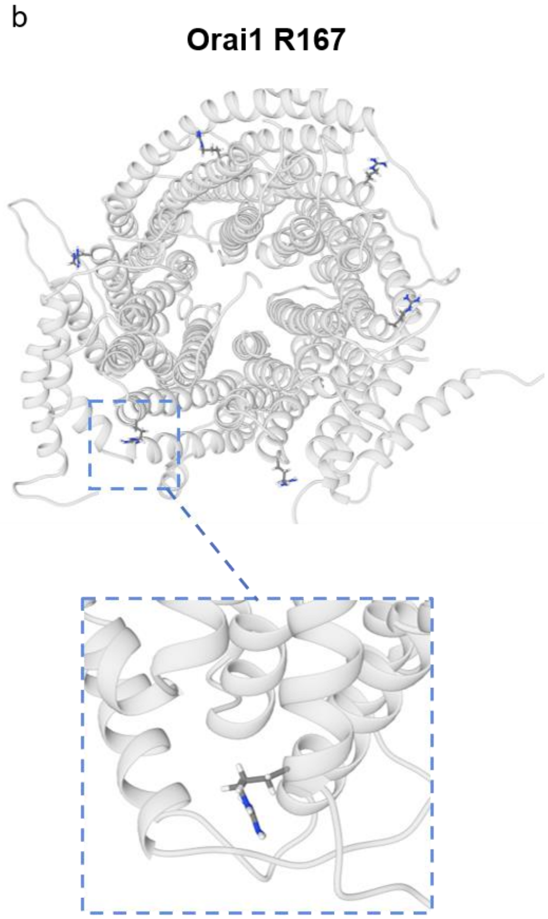


c

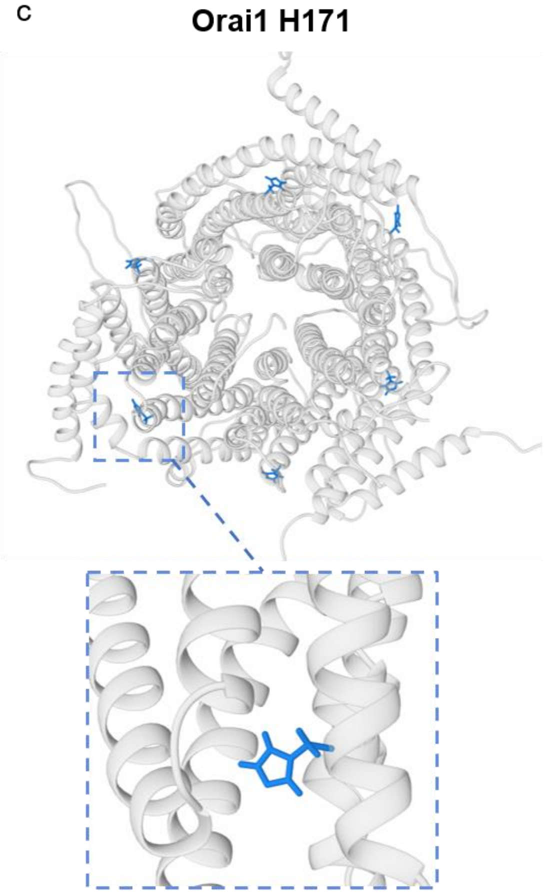

**Supp. Figure 5: Orientation of key residues in the nexus-TM3 interface is critical for UV-induced photocrosslinking**

**a) b) c)** Scheme showing the orientation of a) charged residues K78, E166 and K265, b) R167 and c) H171 in all six subunits of Orai1, taken from our MD simulation on the Orai1 homology model based on dOrai (PDB: 4HKR). The inset a) in red is showing the mean distance between the negatively charged carboxyl oxygen ($COO^-$) of E166 and the amino nitrogen ($NH_3^+$) of K265 and the inset in blue is depicting the mean distance of backbone ($C\alpha$) to backbone between E166 and K265. Mean distances were calculated from respective single distances from all six subunits, listed in the table below; b) is showing a magnified view of R167 in depicted subunit; c) is showing a magnified view of H171 of one respective subunit.

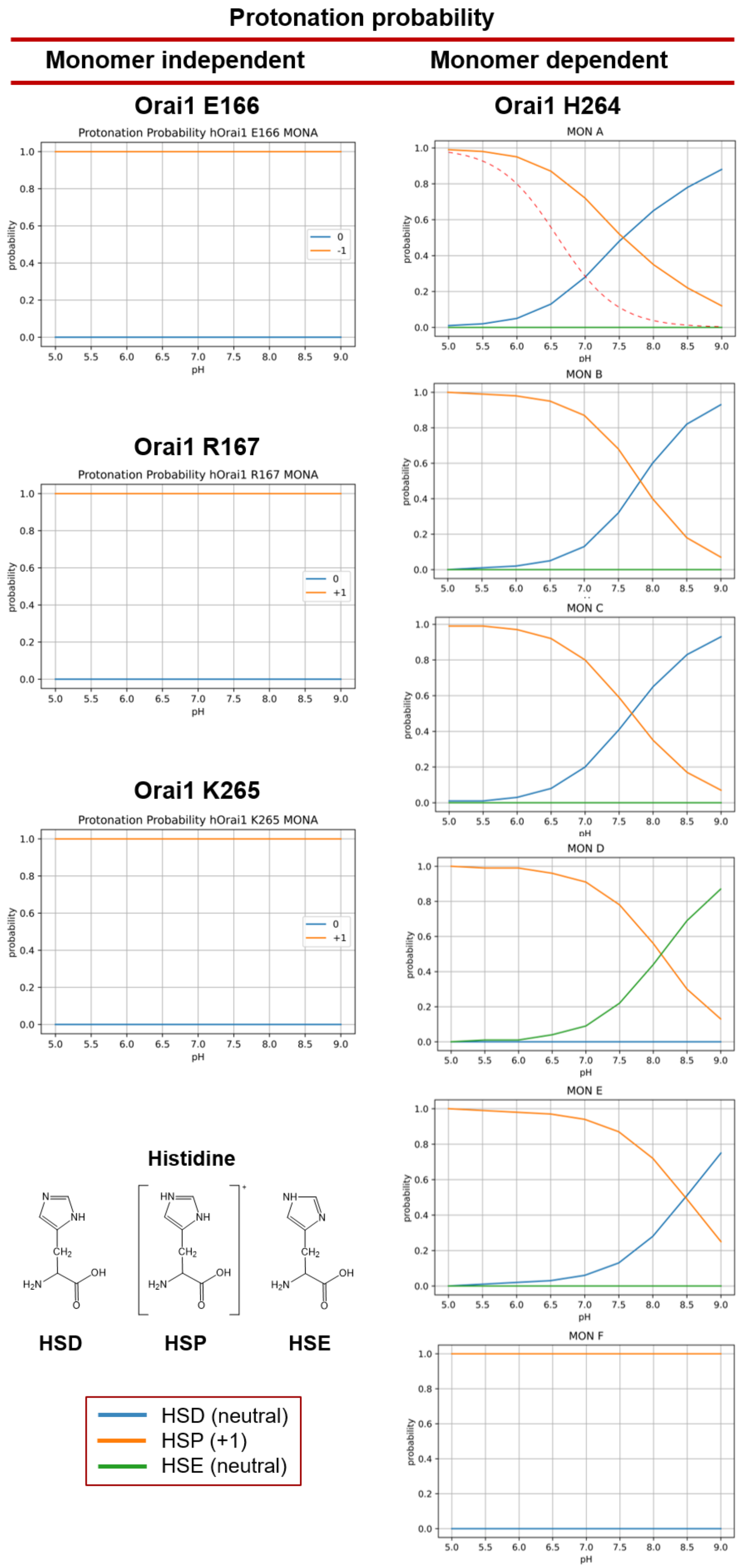
Protonation probability
Monomer independent
Monomer dependent
Orai1 E166
Protonation Probability hOrai1 E166 MONA
Orai1 R167
Protonation Probability hOrai1 R167 MONA
Orai1 K265
Protonation Probability hOrai1 K265 MONA
Orai1 H264
MON A
MON B
MON C
MON D
MON E
MON F
probability
pH
0
-1
+1
Histidine
HSD
HSP
HSE
HSD (neutral)
HSP (+1)
HSE (neutral)

71

**Supp. Figure 6: Titration states of E166, R167, H264 and K265 based on the closed state of Orai1**

Titration curves showing the protonation probability of residues E166, R167, H264 and K265, retrieved by the MC-PBE approach on the homology model of hOrai1 based on dOrai (PDB 4HKR). For Orai1 E166/ R167/ K265 (left), the blue line represents the probability of a neutral state, and the orange line the probability of a charged state. For Orai1 H264 (right), the blue line indicates the probability of the neutral HSD tautomer of histidine, the orange line of the positively charged HSP state, and the green line of the neutral HSE. Chemical structures of HSD, HSP and HSE histidine states are shown in the left corner.

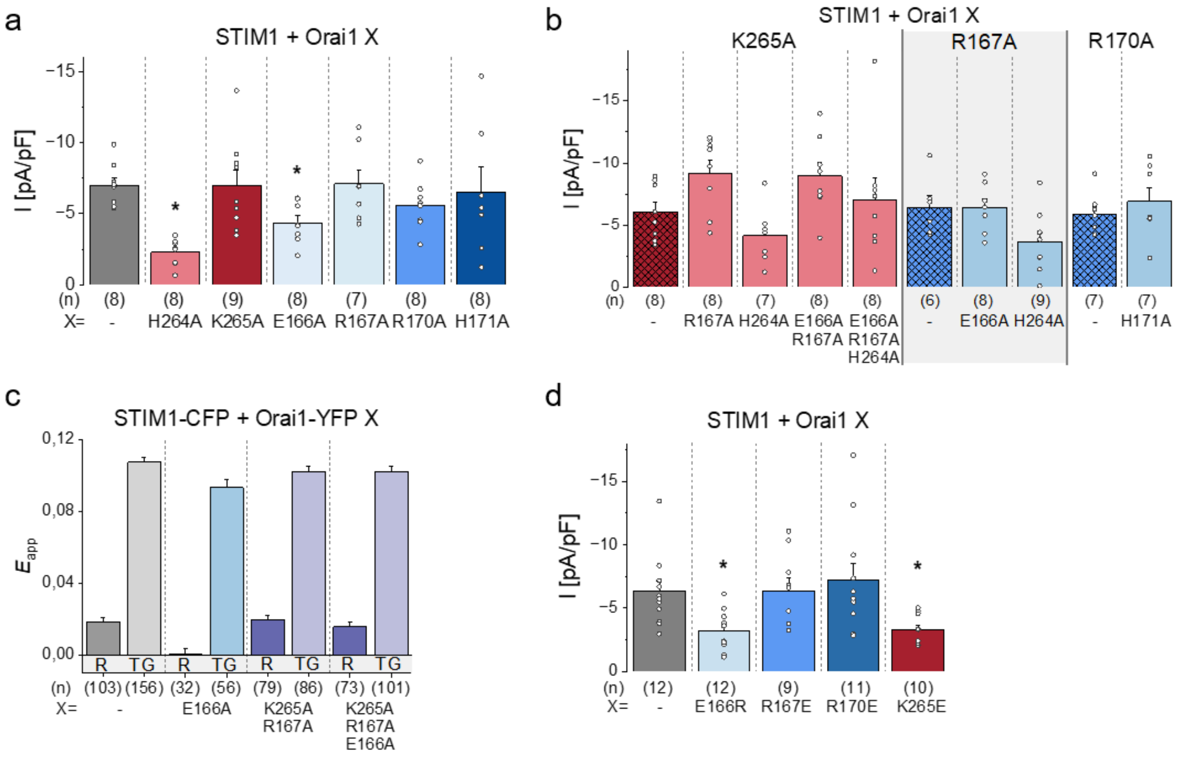


**Supp. Figure 7: Charged residues along the nexus-TM3 interface are required for STIM1-mediated Orai1 channel gating**

**a) b)** Bar diagrams showing maximal current densities of a) Orai1 WT and Orai1 nexus-TM3 alanine-single-mutants and b) Orai1 nexus-TM3 alanine-double/triple-mutants in the presence of STIM1. Statistical significance of Orai1 mutants compared to Orai1 WT are indicated by asterisk * ($p < 0.05$). **c)** Bar diagrams depicting FRET efficiency ($E_{app}$) detecting the interaction of STIM1-CFP with Orai1-YFP WT compared to Orai1 E166A/ K265A R167A/ K265A R167A E166A at resting conditions (R) and after store-depletion with 1 μM Thapsigargin (TG). **d)** Maximal current densities of Orai1 WT and E166R/ R167E/ R170E/ K265E in the presence of STIM1. Single values are indicated as grey circles. Data represent mean values ± SEM of indicated number (n) of experiments.

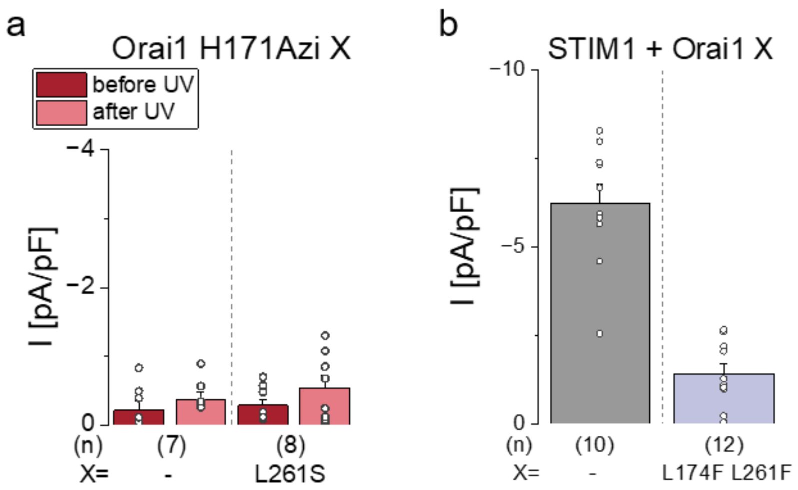


**Supp. Figure 8: Interactions in the upper hydrophobic nexus-TM3 interface are essential for STIM1-mediated Orai1 pore opening**

**a) b)** Bar diagrams showing maximal current densities of a) Orai1 H171Azi and Orai1 H171Azi L261S before and after 15s UV-illumination in the absence of STIM1 and b) Orai1 WT and L147F L261F in the presence of STIM1. Single values are indicated as grey circles. Data represent mean values ± SEM of indicated number (n) of experiments.

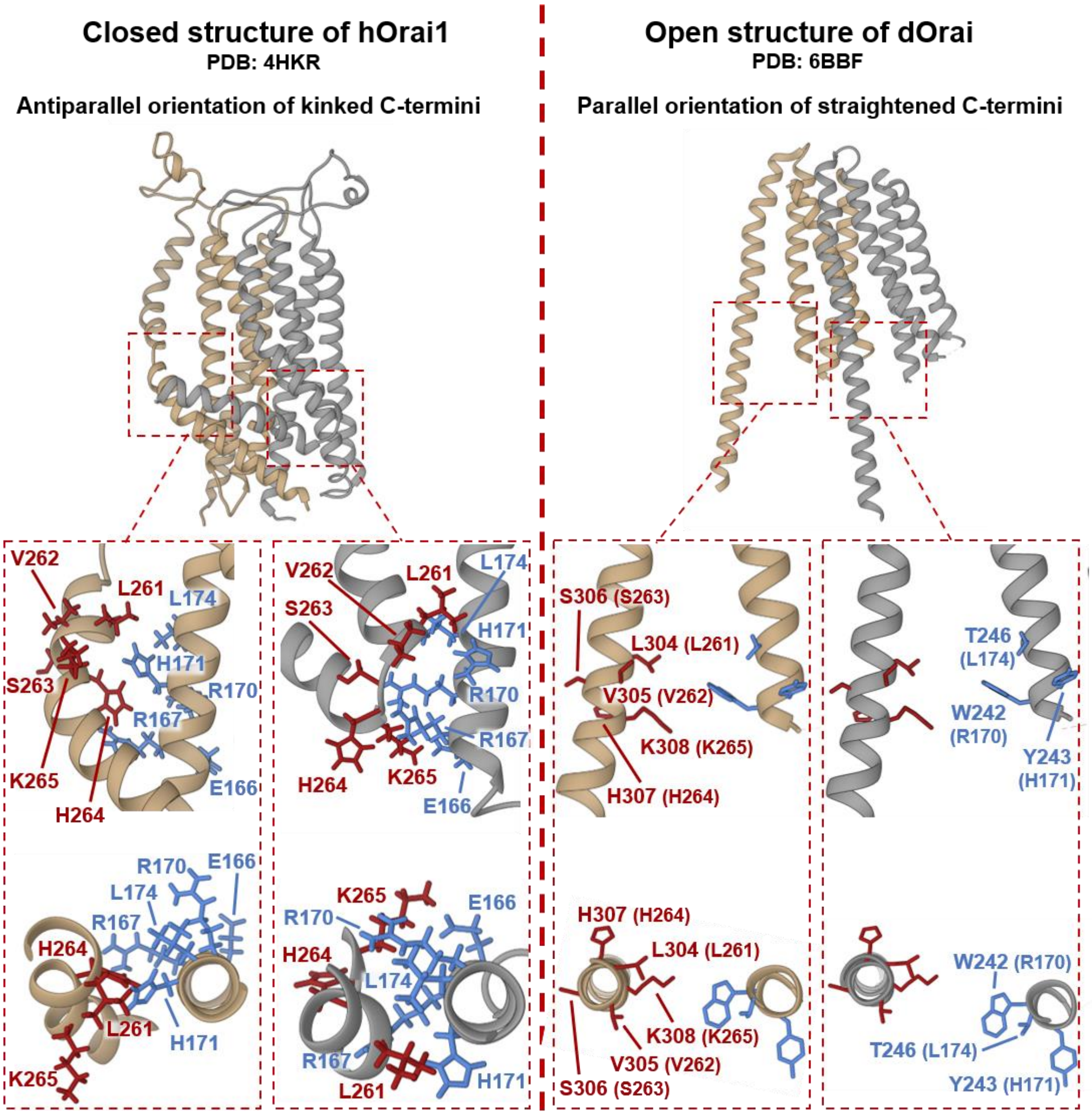


**Supp. Figure 9: The channel conformation of the nexus-TM3 interface of the closed and the open state of Orai1**

Schemes showing a comparison based on the closed structure of the homology model of hOrai1 based on dOrai (PDB: 4HKR) (left) and the open structure of dOrai (PDB: 6BBF) (right), with the insets highlighting the nexus-TM3 interface of two adjacent subunits (brown and grey). The upper panels represent the side view and the lower panels depict the top view of the nexus-TM3 interface. Key residues are highlighted in red (nexus) and blue (TM3). Numbers indicate positions in hOrai1 (left) or dOrai with those of hOrai1 in brackets (right).
